# Regulation of mouse blastocyst primitive endoderm differentiation by p38-mitogen-activated-kinases (p38-MAPKs) is inner-cell-mass (ICM) autonomous and unrelated to cavity expansion defects in ICM specification

**DOI:** 10.1101/2025.09.26.677989

**Authors:** Martina Bohuslavová (née Stiborová), Andrea Hauserová, Rebecca Collier, Joaquin Lilao-Garzón, Silvia Muñoz-Descalo, Alexander W. Bruce

## Abstract

During early mouse blastocyst ICM maturation, we previously described that pharmacological inhibition of p38-MAPK (p38-MAPKi) significantly impairs primitive endoderm (PrE) differentiation from an initially uncommitted population of ICM cells but does not affect pluripotent epiblast (EPI) specification. A recent report details a positive role for blastocyst cavity expansion in assisting ICM lineage formation and marker gene expression. As p38-MAPKi also results in smaller cavity volumes, we addressed to what extent p38-MAPKi mediated impaired PrE differentiation is driven by ICM autonomous or cavity expansion mechanisms. We compared ICM differentiation phenotypes associated with either chemically inhibited cavity volume expansion and p38-MAPKi, on the individual cell and ICM lineage population levels. Whilst recapitulating previously observed decreases in expression of both EPI and PrE markers, we discovered cavity expansion phenotypes are manifest in impaired numbers of specified EPI and increased numbers of uncommitted cells; rather than impaired PrE differentiation, as observed after p38-MAPKi. Moreover, using both 2D ES-cell and 3D ICM organoid models, we show PrE differentiation is also significantly impaired by p38-MAPKi in the absence of a blastocyst cavity; a result recapitulated in cultured immuno-surgically isolated early blastocyst ICMs, in which an outer PrE and inner EPI population are ordinarily formed. These data confirm the early blastocyst requirement for p38-MAPK activity to permit PrE differentiation from uncommitted ICM progenitors is primarily ICM autonomous rather than caused by impaired cavity expansion.

## INTRODUCTION

Mouse preimplantation development forms three blastocyst lineages at implantation (E4.5): an outer, differentiating trophectoderm (TE; placenta precursor; CDX2+) and two inner cell mass (ICM) lineages - the pluripotent epiblast/EPI (foetus progenitors; NANOG+ and SOX2+) and the differentiating primitive endoderm/PrE (a monolayer at the ICM-cavity interface contributing to yolk sac membranes; sequentially activating GATA6, SOX17 and GATA4 expression), as reviewed in (Chazaud and Yamanaka 2016; Plusa and Piliszek 2020). During maturation (E3.5–E4.5), we previously identified an early p38 mitogen-activated-kinase (p38-MAPK) requirement (around E3.5+4 to +7 hours) that permits PrE differentiation from an initially uncommitted ICM population (co-expressing NANOG and GATA6 at E3.5; (Chazaud et al. 2006)), without affecting EPI specification. This phenotype is associated with defects in ribosome-related gene expression, rRNA processing, polysome formation and translation (Thamodaran and Bruce 2016; Bora et al. 2019; Bora et al. 2021a; Bora et al. 2021b). p38-MAPKi is also linked to reduced blastocyst cavity expansion (Bora et al. 2021b). Cavity expansion in mice involves luminal fluid accumulation across the TE driven by ATP1 (N+/K+ ATPase), beginning at the 16-cell stage and coalescing into a single cavity by the 32-cell stage (E3.0–E3.5) (Wiley 1984; Watson 1992; Watson and Barcroft 2001; Motosugi et al. 2005; Ryan et al. 2019; Schliffka et al. 2023).

A previous study used Ouabain (OA) to inhibit ATP1 function and found that impaired blastocyst cavity expansion during maturation is associated with reduced expression of both EPI and PrE marker proteins and with impaired spatial separation of the two lineages within the ICM (Ryan et al. 2019). Building on these findings, we asked to what extent the PrE differentiation defects we previously reported under p38-MAPKi are linked to the observed reduction in blastocyst cavity volume (Bora et al. 2021b), which could reflect compromised TE function, or whether they arise from autonomous ICM mechanisms.

After confirming impaired blastocyst cavity volume expansion with Ouabain (OA), we observed a concomitant and consistent reduction in the expression of EPI markers (NANOG and SOX2) and PrE markers (GATA6, SOX17 and GATA4) in ICM cells. Additionally, by the late-blastocyst stage (E4.5), OA-treated ICMs contained significantly fewer specified EPI cells than controls, while the number of specified/differentiating PrE cells was unchanged. The reduced EPI cell numbers were counterbalanced by an increased population of uncommitted ICM cells co-expressing NANOG and GATA6, indicating impaired EPI specification. These data reveal that EPI specification, rather than PrE differentiation, is more sensitive to impaired cavity expansion.

Moreover, the OA-induced phenotype differs from the inverse PrE-defect observed under p38-MAPKi conditions during the same developmental window, supporting the role of active p38-MAPK signalling in promoting PrE differentiation at the ICM level, independently of cavity expansion. Substantiating this conclusion, outer PrE cell formation in immuno-surgically isolated (IS) ICMs of early blastocysts lacking TE and a cavity is impaired after p38-MAPKi; and PrE-like differentiation is also reduced after p38-MAPKi in employed 2D ES-cell and 3D ICM organoid models (Schroter et al. 2015; Mathew et al. 2019). Collectively, these findings reinforce the role of p38-MAPK signalling in PrE differentiation and highlight the distinct but interacting contributions of cavity volume expansion and ICM-autonomous mechanisms to blastocyst cell fate.

## RESULTS

### Ouabain (OA) and p38-MAPKi induced blastocyst cavity expansion defects result in distinct ICM cell fate population phenotypes

We previously reported that cultured mouse blastocysts (E3.5-E4.5) exposed to p38-MAPKi (using SB220025 (Jackson et al. 1998)) show more uncommitted ICM cells (NANOG+ & GATA6+) and fewer differentiating PrE cells (GATA4+), with little effect on EPI specification (NANOG+ but lacking GATA6/GATA4); indicating impaired PrE progenitor differentiation (Thamodaran and Bruce 2016). Additionally, p38-MAPKi–treated blastocysts have significantly reduced cavity volume (Bora et al. 2021b). Other studies show that experimentally reducing cavity expansion (via OA-mediated ATP1 inhibition or cavity-fluid removal) lowers expression of EPI and PrE marker proteins (SOX2 and GATA4) and impairs ICM cell sorting (Ryan et al. 2019). Therefore, we aimed to extend these findings and determine how OA treatment affects the derivation and final EPI versus PrE composition of the late-blastocyst (E4.5) ICM lineages. We also sought to compare OA-phenotypes with those of p38-MAPKi during the same maturation window to understand whether p38-MAPKi–mediated impairment of PrE specification/differentiation arises from cavity expansion effects *per se* or from ICM-autonomous mechanisms.

We first recapitulated our published p38-MAPKi blastocyst maturation phenotypes (E3.5–E4.5), showing impaired PrE specification/differentiation and reduced cavity volume expansion (Thamodaran and Bruce 2016; Bora et al. 2021b), assessed by immunofluorescence (IF) staining for GATA4/GATA6 and NANOG or GATA4/GATA6 and SOX2 (Figs. S1–S6; indicating ICM cell fate resolution, quantified marker protein expression and blastocyst cavity volume measurements). Following an analogous approach, we cultured early blastocysts (E3.5) in the presence of OA (as previously described (Ryan et al. 2019)) or DMSO vehicle control for 12 hours (E4.0) or 24 hours (E4.5). Consistent with previous reports (Manejwala et al. 1989; Bagnat et al. 2007), OA treatment caused significant cavity-volume defects and impaired blastocyst hatching at E4.5, without affecting total embryo or ICM cell numbers at either time point (Fig. 1).

**Fig. 1:**
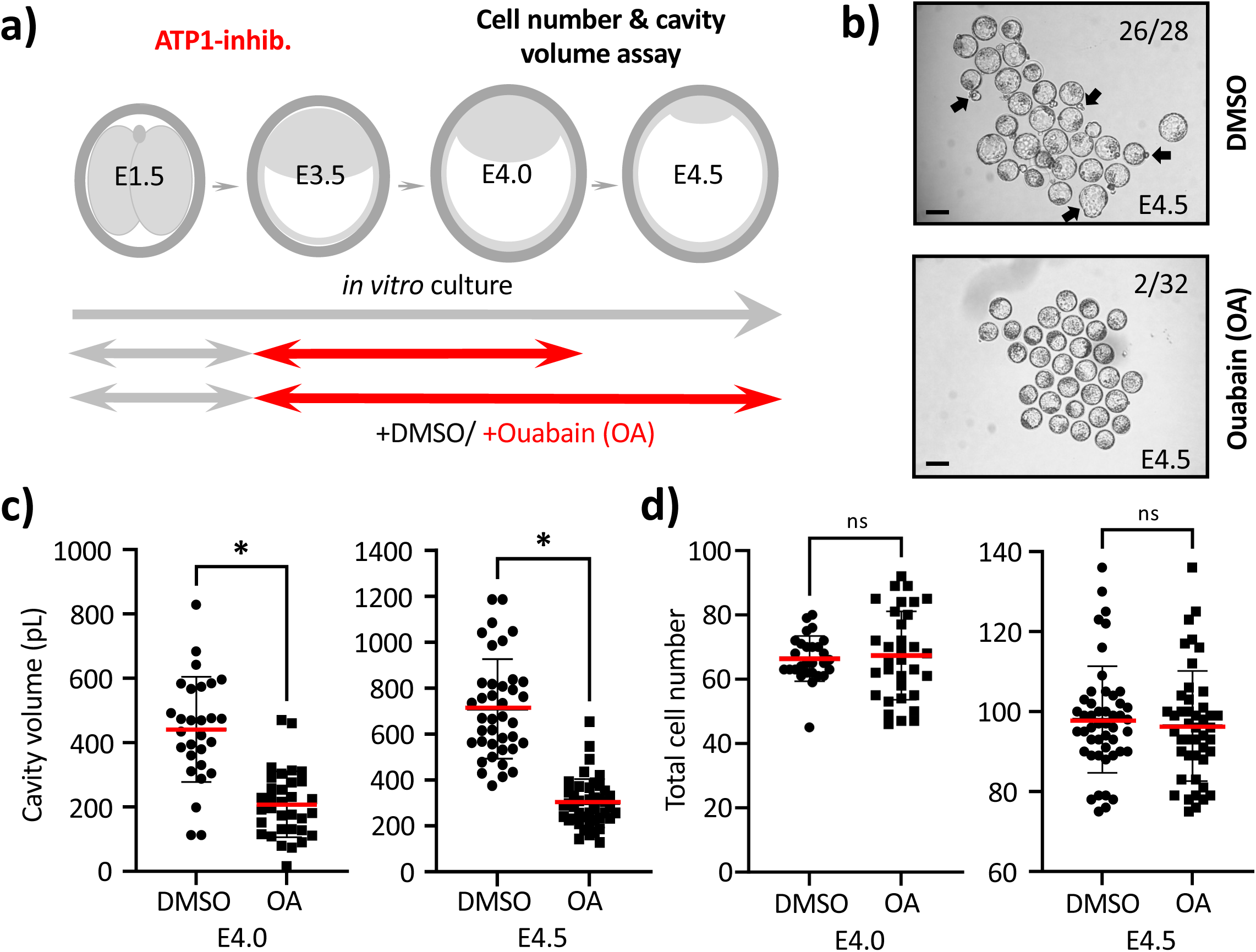
Ouabain treatment during blastocyst maturation causes reduced cavity size and impaired hatching without affecting total embryo cell number: **a)** Experimental scheme of Ouabain (OA) treatment and assay points during blastocyst maturation period (E3.5-E4.5). **b)** Bright-field micrographs of control (DMSO - upper) or OA-treated (lower) blastocyst groups at E4.5; note impaired blastocyst hatching (see arrows in control) after OA treatment (scale bar - 100μm). **c)** Average total cell number per control (DMSO) or OA-treated blastocyst group at E4.0 (left) or E4.5 (right). **d)** Average cavity volume per control or OA-treated blastocyst group at E4.0 (left) or E4.5 (right). Panels c) and d); arithmetic means (red bars), standard deviations and statistically significant differences are highlighted (for normal or non-normally distributed data, unpaired t-test or Mann-Whitney tests were employed, respectively. *p<0.05; see supplementary data/stats Excel tables).

To extend these findings, we conducted confocal microscopy IF imaging to quantify: (i) ICM population sizes of specified EPI, differentiating PrE, and uncommitted progenitors (unique to this study), and (ii) the expression levels of ICM lineage markers (as conducted in Ryan et al. 2019)). This served two purposes. Firstly, to corroborate and expand the OA-related ICM cell-fate data, and secondly, to provide an OA-related reference dataset for comparing phenotypes with those induced by p38-MAPKi. Furthermore, to broaden the repertoire of lineage markers and developmental timepoints analysed under OA-treatment conditions, we probed combinations of EPI markers (NANOG or SOX2) with PrE markers (GATA6, SOX17, or GATA4) following OA treatment. Under ±OA conditions, we first determined the average number of cells contributing to ICM sub-populations at E4.0, based on assigning combinations of expressed lineage markers: NANOG, GATA6, and GATA4. These are summarised as population percentage contributions across the whole ICM (Fig. 2) or as absolute cell numbers per embryo (Fig. S7) with associated per cell quantified protein expression levels (Fig. S8). At E4.0, many ICM cells in both control (DMSO) and OA-treated blastocysts remained NANOG+ & GATA6+ (N+G6+), i.e., uncommitted progenitors, indicating incomplete resolution of EPI and PrE-specified progenitors by this stage (yellow arrows in Fig. 2b). However, OA increased the percentage of uncommitted ICM cells compared with DMSO (57.0% vs 47.0%, Fig. 2c), and reduced the fraction of EPI-specified cells (N+G6−) from 20.2% in DMSO to 14.4% in OA. The proportion of PrE-specifying cells (N−G6+) showed a non-significant difference between groups (DMSO 32.8% vs OA 28.6%, Fig. 2c). The fraction of ICM cells expressing the later PrE marker GATA4 was also not significantly different between groups (DMSO 17.6% vs OA 17.0%, Fig. 2c). Since OA did not affect total blastocyst cell number (Fig. 1), the shifts in ICM composition at E4.0 with OA are also reflected in the overall cell-number changes, with fewer EPI-specified cells (N+G6−) and more uncommitted cells (N+G6); and, PrE-specified cell numbers, marked by GATA6 in the absence of NANOG or by GATA4, did not significantly differ (Fig. S7). Although OA-treatment selectively impaired EPI specification, it did not alter the average per-cell NANOG expression within EPI (N+G6−) or uncommitted (N+G6+) cells. By contrast, GATA6 expression was significantly reduced in both uncommitted (N+G6+) and specifying PrE (N−G6+) cells, even though such PrE-related populations were unaffected in size. Notably, GATA6 expression remained unchanged in those PrE cells that had already begun to express GATA4 (N−G6+G4+), and GATA4 expression levels themselves were unchanged (Fig. S8). Taken together, OA-induced cavity-size reduction at E4.0 correlates with impaired EPI progenitor specification from the initially uncommitted E3.5 ICM, without altering average NANOG levels in those cells, but it does not impair PrE specification/differentiation, even though GATA6 is reduced in some uncommitted and early PrE subpopulations that have not yet expressed GATA4.

**Fig. 2:**
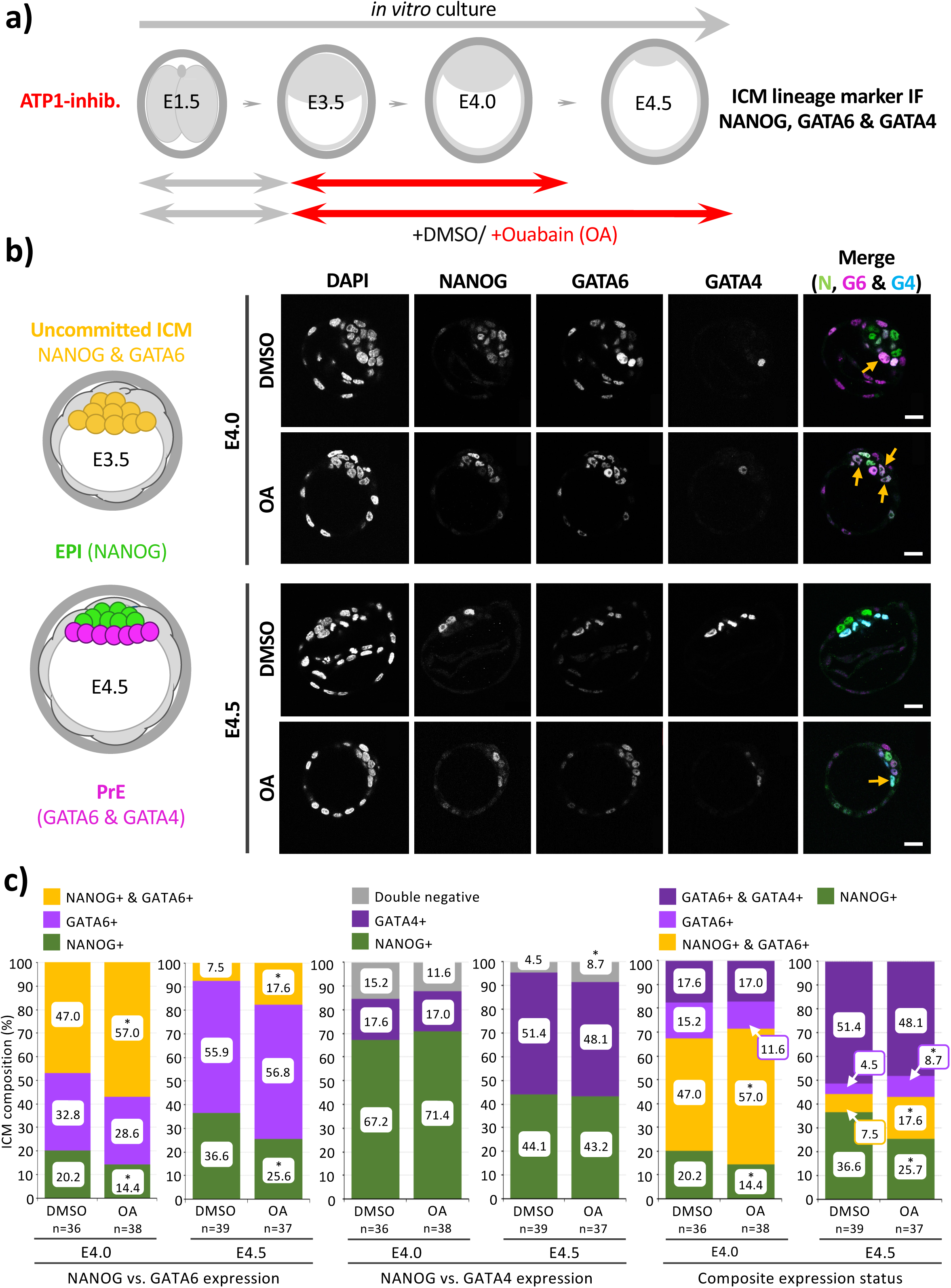
Ouabain treatment during blastocyst maturation impairs EPI specification from uncommitted ICM progenitors without affecting PrE differentiation (IF: NANOG/GATA6 & GATA4): **a)** Experimental scheme of Ouabain (OA) treatment and assay points during blastocyst maturation period (E3.5-E4.5). **b)** Left - Schematic of typical expression of ICM lineage marker proteins in uncommitted early-blastocyst (E3.5) stage (NANOG & GATA6 co-expression - yellow) and at the late-blastocyst (E4.5) stage (EPI – NANOG alone – green, PrE – GATA6 & GATA4 in the absence of NANOG - magenta). Right - Exemplar confocal single z-section micrographs of control (DMSO) or OA-treated blastocysts IF assayed for combined NANOG, GATA6 & GATA4 expression at E4.0 & E4.5 (scale bar - 20μm). **c)** Average cellular percentage ICM composition of control (DMSO) or OA-treated blastocysts IF assayed for cell lineage marker expression at E4.0 & E4.5; delineated as i. uncommitted (co-expressing NANOG & GATA6), PrE (expressing GATA6 without NANOG) and EPI (expressing NANOG without GATA6) cells (left panel), ii. PrE (expressing GATA4 without NANOG) and EPI (expressing NANOG without GATA4) cells (central panel), or iii. uncommitted (co-expressing NANOG & GATA6), PrE (expressing GATA6 alone or GATA6 plus GATA4 without NANOG) and EPI (expressing NANOG without GATA6 or GATA4) cells (right panel). Statistically significant differences highlighted (Z-test; *p<0.05, see supplementary data/stats Excel tables; embryo n-numbers highlighted). *See Fig. S7 for average total cell number data/ per assayed embryo.* Note, OA-treatment results in unaffected PrE specification, impaired EPI formation and increased incidence of uncommitted; see yellow arrows in panel b).

We next asked what effect OA treatment would have on E4.5 stage blastocyst ICMs (Fig. 2c and Fig. S7). A smaller population of uncommitted ICM progenitors (N+/G6+) remained observable in both DMSO and OA-treated embryos, reflecting ongoing ICM fate resolution during later blastocyst maturation. However, as at E4.0, this uncommitted population was significantly greater with OA treatment than in control DMSO conditions (7.5% in DMSO vs 17.6% in OA of the total ICM). Similarly, there was a significant reduction in the proportion of specified EPI cells (N+/G6−) in OA-treated embryos (DMSO 36.6% vs OA 25.6%), while the contribution of PrE-specified cells (N−G6) did not significantly differ between groups (DMSO 55.9% vs OA 56.8%). There was also no significant difference in the proportion of ICM cells expressing the later PrE marker GATA4 (N−G4+; DMSO 51.4% vs OA 48.1% - Fig. 2c). Thus, OA treatment throughout the entire blastocyst maturation period, which also impairs cavity expansion (Fig. 1), uniquely disrupts the specification of EPI progenitors from initially uncommitted ICM cells, while PrE specification is not similarly affected (these phenotypes are also evident when looking at the averaged total ICM cell numbers at E4.5, Fig. S7). This indicates that the mid-blastocyst (E4.0) specification defects persist through to the late-peri-implantation stage. An analysis of the average expression of levels of all three lineage markers showed that OA treatment significantly reduced NANOG expression in both EPI-specified cells (N+G6−) and uncommitted cells (N+G6+). GATA6 and GATA4 expression was also reduced in PrE-specified and differentiating cells (i.e. N−G6+ and N−G6+G4+), but GATA6 expression in uncommitted cells (N+G6+) remained unaffected (Fig. S8). These findings align with prior reports that OA treatment diminishes EPI (SOX2) and PrE (GATA4) marker expression up to E4.0, even though those earlier studies examined blastocysts flushed at E3.5 (Ryan et al. 2019) rather than those cultured from the 2-cell stage.

We repeated both OA treatment regimens and assayed SOX2 (as in (Ryan et al. 2019)) and SOX17 expression as alternative markers for the EPI (SOX2 (Avilion et al. 2003; Wicklow et al. 2014)) and PrE (SOX17 (Niakan et al. 2010; Artus et al. 2011)) lineages (Fig. 3 and Fig. S9). At E4.0, OA caused a significant reduction in the proportion of EPI-specified cells that express only SOX2 (S2+S17−; DMSO 51.1% vs OA 38.9%), while the proportion of PrE-specified cells (S2−S17+) did not differ significantly (DMSO 20.3% vs OA 16.4%). OA also led to a significantly higher fraction of ICM cells co-expressing both markers (S2+S17+; DMSO 28.6% vs OA 44.7%). These trends were mirrored in the averaged total ICM cell numbers (Fig. S9). Interestingly, despite SOX17 being a temporally later PrE marker than GATA6 (in the absence of NANOG), a large proportion of mid-blastocyst (E4.0) ICM cells, even in control DMSO conditions, initiated SOX17 expression while SOX2 remained present. Moreover, such co-expression of SOX2 and SOX17 was further enhanced by OA treatment. Although SOX2/SOX17 co-expression resembles uncommitted cells co-expressing NANOG/GATA6, it may be more appropriate to interpret this state as a transient, conflicted phase of ICM cell-lineage specification, arising during the transition from the early-blastocyst stage rather than a maintained condition; given early- (E3.5) stage blastocysts express GATA6 but not SOX17(Chazaud et al. 2006; Niakan et al. 2010; Thamodaran and Bruce 2016). Furthermore, whilst ICM cells of maturing blastocysts ordinarily transit this state, it is potentiated by OA treatment (and it is marked by enhanced SOX2, yet unchanged SOX17, expression levels – Fig. S10) and associated cavity volume expansion defects (Fig. 1). Despite this, when OA treatment is applied from E3.5 to E4.5, the percentage compositions of ICM cells defined by SOX2 and SOX17 expression/co-expression was not statistically different (Fig. 3). This resolution of ICM cell fate contrasts with the results of the NANOG/GATA6 analysis, where EPI specification from uncommitted ICM progenitors is impaired at E4.0 and persists through E4.5 (Fig. 2 and Fig. S7). The discrepancy may reflect differential sensitivity and mechanosensory regulation of individual lineage-marker proteins in response to impaired cavity expansion (Fig. 1). Notably, by E4.5, the per-ICM-cell expression levels of both SOX2 and SOX17 (where detected) are significantly reduced across all expression/co-expression categories (Fig. S10), consistent with prior data showing OA-induced reductions in EPI (SOX2) and PrE (GATA4) markers after OA treatment between E3.5 and E4.0 (Ryan et al. 2019). Collectively, these findings confirm that OA-mediated reductions in cavity size (Fig. 1) are associated with reduced expression of markers for both ICM lineages, but preferentially impede the differentiation of EPI progenitor cells rather than their PrE counterparts (Figs. 2–3 and Figs. S7–S10).

**Fig. 3:**
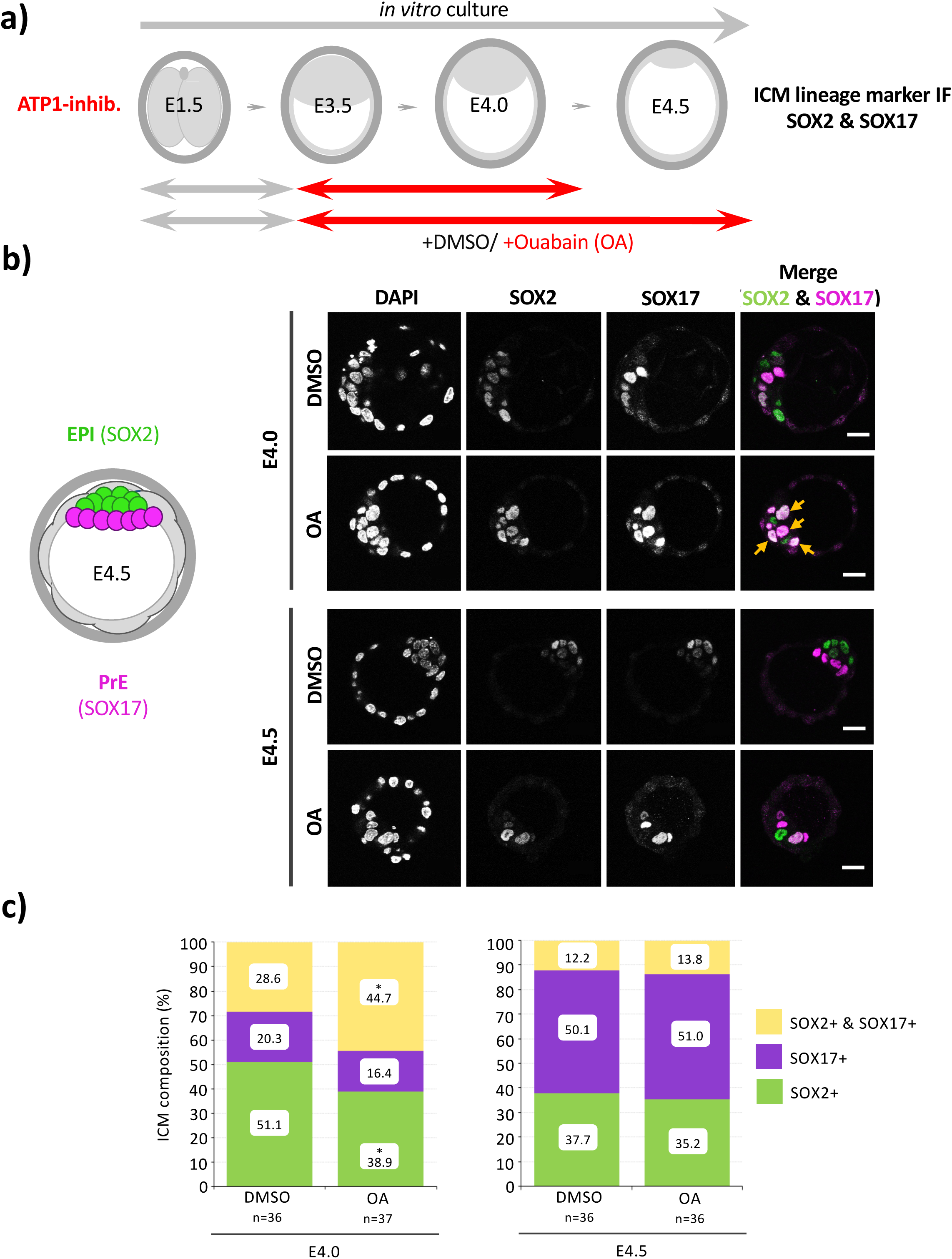
Ouabain treatment during blastocyst maturation impairs EPI specification without affecting PrE differentiation (IF: SOX2 & SOX17): **a)** Experimental scheme of Ouabain (OA) treatment and assay points during blastocyst maturation period (E3.5-E4.5). **b)** Left -Schematic detailing typical expression of ICM lineage marker proteins at the late-blastocyst (E4.5) stage (EPI – SOX2 alone – green, PrE – SOX17 - magenta). Right - Exemplar confocal single z-section micrographs of control (DMSO) or OA-treated blastocysts IF assayed for combined SOX2 & SOX17 expression at E4.0 & E4.5 (scale bar - 20μm). **c)** Average cellular percentage ICM composition of control (DMSO) or OA-treated blastocysts at E4.0 (left) & E4.5 (right); delineated as co-expressing (SOX2 & SOX17), PrE (expressing SOX17 without SOX2) and EPI (expressing SOX2 without SOX17) cells. Statistically significant differences highlighted (Z-test; *p<0.05, see supplementary data/stats Excel tables; embryo n-numbers highlighted). *See Fig. S9 for average cell number population data/ per assayed embryo.* Note, OA-treatment results in unaffected ICM specification of PrE at E4.0 & E4.5 and impaired formation of EPI and increased incidence of co-expressing cells at E4.0; see orange arrows in panel b).

In summary, these analyses show that OA-induced impairment of blastocyst cavity volume expansion affects the appropriate formation of ICM subpopulations by impairing EPI specification while leaving PrE specification relatively unaffected (Figs. 1-3). This is accompanied by reduced expression of both EPI and PrE marker proteins (Figs. S8 & S10), supporting and expanding a mechanosensory role for cavity expansion in maintaining ICM marker protein expression, as previously reported (Ryan et al. 2019). This phenotype, however, contrasts with our earlier p38-MAPKi results (Thamodaran and Bruce 2016; Bora et al. 2019; Bora et al. 2021b), where PrE specification/differentiation from uncommitted ICM progenitors is compromised rather than EPI specification (Figs. S1-S6). Thus, although p38-MAPKi inhibition is associated with reduced cavity expansion, it is unlikely to be a major driver of the PrE-specific phenotype previously observed.

### PrE differentiation in early blastocyst (E3.5) isolated ICMs is impaired by p38-MAPKi

To address the hypothesis p38-MAPK supports PrE specification/differentiation via an ICM autonomous mechanism, we assayed ICM lineage formation in a context devoid of TE and associated cavity expansion, using immuno-surgically (IS) isolated ICMs. Depending on exact timing, cultured IS-derived early ICMs from E3.5 mouse blastocysts, where outer TE cells are removed by complement-mediated lysis (Fig. S11; (Solter and Knowles 1975)), either reform an outer TE layer (sometimes reconstituting a cavity) or, if TE fate has already irreversibly committed (Posfai et al. 2017), develop an outer differentiated PrE encapsulating NANOG-expressing EPI cells (Wigger et al. 2017). We first reproduced these findings: ICMs isolated around 91 hours post-hCG (relative to female superovulation) and cultured for 24 hours reconstituted a TE expressing CDX2, whereas those isolated at 96 hours formed an outer PrE layer marked by GATA4 surrounding NANOG-expressing EPI cells (Fig. S11). Using the 96-hour timepoint, we identified a working concentration of 5 μM SB220025 (p38-MAPKi) that did not cause ICM cytotoxicity after 24 hours in culture (Fig. S12); despite greater cytotoxic sensitivity of IS-isolated ICMs compared to p38-MAPKi in intact blastocysts, a similar 5μM (versus 20μM) treatment in E3.5–E4.5 cultured blastocysts still produced a small but significant increase in the proportion/ number of uncommitted cells (compare Figs. S13–S14 with previous results (Thamodaran and Bruce 2016) or those recapitulated here in Figs. S1–S6). The increased p38-MAPKi sensitivity of isolated ICMs most probably reflects enhanced availability of the inhibitor to ICM cells, in a context lacking overlying TE cells in blastocyst treatments.

Given our previous finding of a narrow and irreversible minimal window for p38-MAPKi–mediated PrE impairment before the mid-blastocyst stage (i.e. between E3.5 and E4.0; (Bora et al. 2021b)), we tested the effect of 12 hours of ±p38-MAPKi (5 μM SB220025) followed by 12 hours in standard media (to reach the equivalent E4.5 stage) on ICMs isolated at the 96-hour post-hCG timepoint, assessing EPI (SOX2) and PrE (GATA4) formation (Fig. 4 and Fig. S15). We found that, in accord with similar treatment of intact blastocysts (E3.5-E4.5), p38-MAPKi efficiently impaired specification and differentiation of the PrE (marked by GATA4 expression in the absence of SOX2), in stark contrast to DMSO vehicle-treated controls exhibiting a surface PrE layer. Indeed, we rarely observed cells positive for GATA4 expression in any IS-isolated ICMs after p38-MAPKi treatment, whereas all cells expressed the EPI marker SOX2. However, we did observe that although virtually all ICM cells expressed SOX2 after p38-MAPKi treatment, the average per-cell SOX2 expression was significantly reduced compared with controls (Fig. S15). Consistently, this p38-MAPKi-induced reduction in SOX2 expression (in the absence of GATA6/GATA4 expression) was also seen in intact blastocysts (Fig. S6), although the trend did not extend to the alternative EPI marker NANOG (Fig. S3); indicating differing relative stabilities in EPI lineage marker expression under p38-MAPKi conditions. Overall, we interpret these data as indicating PrE specification and differentiation requires active p38-MAPK signalling, even in a TE- and cavity-free context, in a manner consistent with an ICM-autonomous mechanism of cell-fate resolution.

**Fig. 4:**
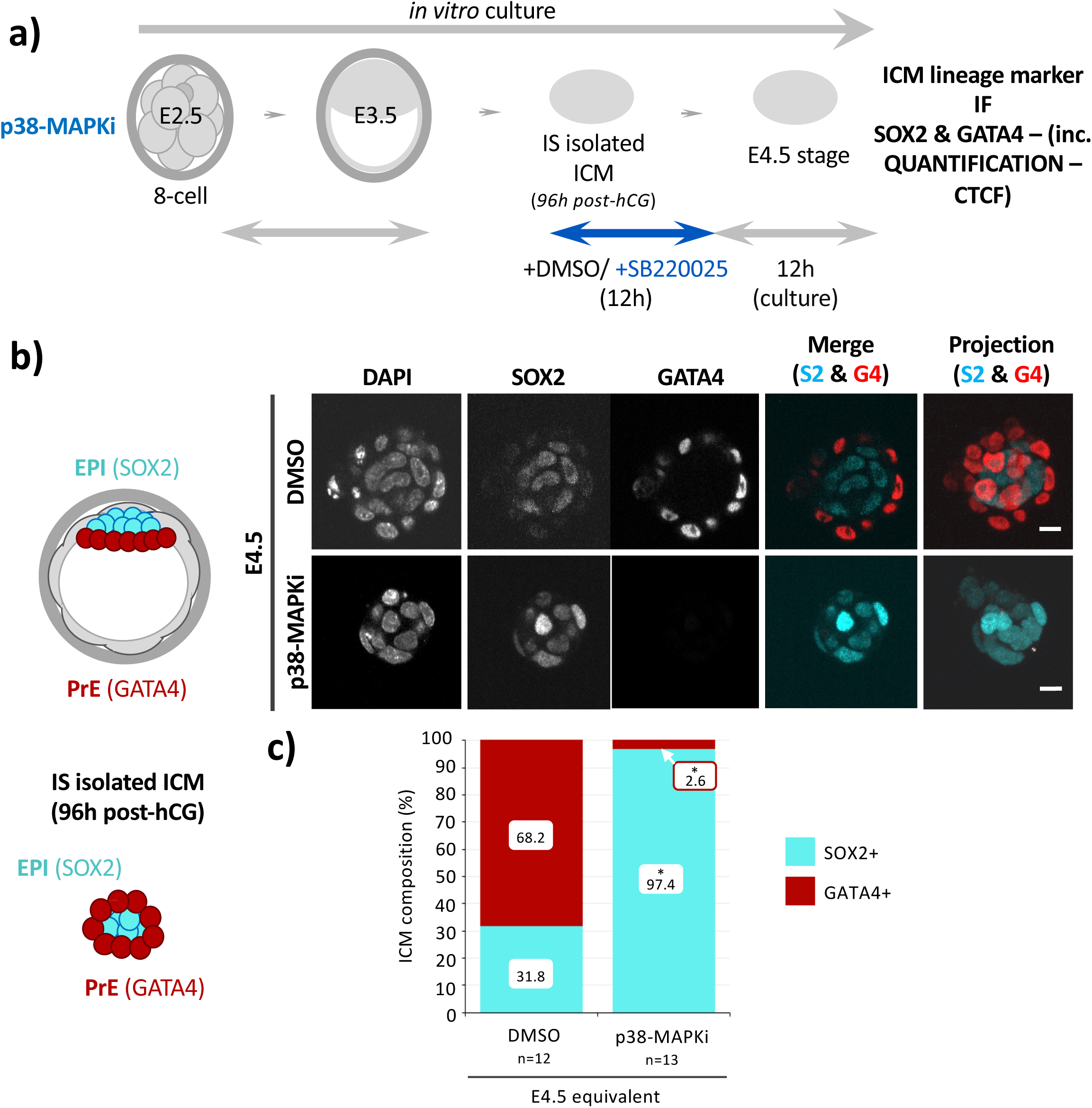
p38-MAPKi treatment of early-blastocyst (E3.5) stage isolated ICMs effectively impairs PrE lineage differentiation: **a)** Experimental scheme of early-blastocyst (E3.5 – 96 hours post-hCG superovulation microinjection) stage ICM isolation and 12-hour culture under control (DMSO) or p38-MAPKi (5μM SB220025) conditions before transfer into regular growth media for another 12 hours, until the equivalent late-blastocyst (E4.5) stage, and IF staining for ICM lineage markers (SOX2 & GATA4). **b)** Left - Schematic detailing typical expression of ICM lineage marker proteins at the late-blastocyst (E4.5) stage or in isolated and cultured early-blastocyst stage ICMs (i.e. after TE commitment) (EPI – SOX2 alone – cyan and PrE – red, in the absence SOX2 expression). Right - Exemplar confocal single z-section micrographs of control (DMSO) or p38-MAPKi treated ICMs IF stained for combined SOX2 & GATA4 expression (scale bar - 20μm); plus, merged images and a maximally projected z-section image (right-most micrographs). **c)** Average cellular percentage ICM composition of control (DMSO) or p38-MAPKi treated ICMs; delineated as EPI (SOX2) or PrE (GATA4) expressing cells. Statistically significant differences highlighted (unpaired t-test; *p<0.05, see supplementary data/stats Excel tables; ICM n-numbers shown). *See Fig. S15 for average cell number population data/ per assayed isolated ICM*.

### PrE differentiation in inducible 2D ES-cell and 3D ES-cell ICM organoid models is sensitive to p38-MAPKi

Previous work with the transgenic 2D mouse ES-cell line *Tet::GATA6mCherry* (Schroter et al. 2015; Mathew et al. 2019) shows that a 6-hour transient exposure to doxycycline (+DOX) induces co-expression of endogenous NANOG and the recombinant, DOX-induced GATA6-mCherry protein. This co-expression mirrors the uncommitted state of early blastocyst ICM (Chazaud et al. 2006). Furthermore, after 48 hours of *in vitro* culture without DOX (-DOX), and even under conditions that promote ES cell pluripotency (i.e., +serum +Leukaemia Inhibitory Factor/LIF), cells adopt mutually exclusive patterns of NANOG and GATA6 expressing populations. In this model, GATA6 expression is accompanied by induced expression of other PrE markers such as SOX17 and GATA4 (Schroter et al. 2015; Mathew et al. 2019). Therefore, we adopted this ES-cell–based ICM cell fate model to test the effect of p38-MAPKi, investigating whether p38-MAPK promotes PrE fate independently of cavity-expansion–related morphologies, acting instead at the cell (and by inference the ICM) population level.

Accordingly, we exposed *Tet::GATA6mCherry* ES cells to either control DMSO or increasing concentrations of SB220025/p38-MAPKi (5, 10, and 20μM) during the 6-hour +DOX induction period and immediately assayed for endogenous NANOG and GATA4 and recombinant GATA6-mCherry expression by confocal-IF microscopy (Figs. S16 & S17). Under DMSO control conditions, we confirmed co-expression of NANOG and GATA6-mCherry in the absence of detectable GATA4 (N+G6+) in 71.7% of assayed cells, resembling the uncommitted early-blastocyst ICM state and confirming successful GATA6-mCherry induction. Interestingly, 21.5% of these DMSO-treated cells showed downregulated NANOG (N−G6+), suggesting initiation of differentiation toward a PrE-like fate. In all p38-MAPKi treatment conditions, the proportion of PrE-like cells (N−G6+) was significantly reduced, and the proportion of uncommitted cells (N+G6+) was significantly increased. This effect was most pronounced using 10 μM SB220025/p38-MAPKi (N+G6+ = 93.3% and N−G6+ = 2.7%), which, unlike the 20 μM treatment, did not show cytotoxicity. Additionally, quantification of marker protein expression revealed significantly higher cellular NANOG levels in the 10 μM p38-MAPKi-treated uncommitted (N+G6+) cells compared to DMSO controls (Fig. S18); suggesting the reduced derivation of PrE-like cells (N−G6+) is antagonised by p38-MAPKi through enhanced induction of endogenous NANOG expression.

We next tested the effect of p38-MAPKi treatment, applied during the 6-hour +DOX induction period, on the formation of the previously described ICM-like lineages after a further 24 hours of culture in normal media without DOX; assaying endogenous NANOG, GATA6, and GATA4 expression (Fig. 5). We found that p38-MAPKi-treated ES cells were significantly impaired in generating PrE-like lineages that lack NANOG but express GATA6 (N−G6+): 14.4% in DMSO controls versus 4.3% with p38-MAPKi. Similarly, PrE-like cells expressing GATA4 (N−G4+) were reduced: 14.8% in controls versus 4.5% after p38-MAPKi. This impairment was accompanied by a significant shift toward uncommitted lineages that co-express NANOG and GATA6 (N+G6+): 30.6% in controls versus 42.8% with p38-MAPKi. The proportion of EPI-like cells expressing only NANOG (N+G6−) was not significantly altered (45.2% in controls vs 44.0% in p38-MAPKi); however, there was a significant increase in cells expressing NANOG in the absence of GATA4 (N+G4−), indicative of a population comprising both EPI-like cells and uncommitted cells (that would also express GATA6). Collectively, these results follow those obtained after p38-MAPKi in intact blastocysts (Thamodaran and Bruce 2016; Bora et al. 2019; Bora et al. 2021b) (recapitulated here; Figs. S1-S3) and IS-isolated ICMs (Fig. 4 and Fig. S15) cultured through the blastocyst maturation period (E3.5–E4.5). Unlike the timepoint immediately after +DOX induction, we did not observe a significant increase in NANOG expression in cells in the uncommitted state (N+G6+), although GATA6 expression was significantly elevated, including in the subset of cells that had initiated PrE differentiation (N−G6+). NANOG expression was also significantly higher in cells that progressed toward an EPI-like state (N+G6− and N+G4−; Fig. S19). When considering only NANOG and GATA4 status, p38-MAPKi caused a significant increase in the proportion of otherwise uncommon cell populations co-expressing NANOG and GATA4 (N+G4+: 2.0% with DMSO vs. 8.9% after p38-MAPKi), further indicating PrE-specification and differentiation defects. Indeed, aggregating all possible marker combinations by their ICM expression status at the late (E4.5) blastocyst stage as PrE-like, EPI-like, ICM-like, or triple negative (where any co-expression of NANOG and/or GATA6 and GATA4 is classified as uncommitted/ICM-like), reveals that only PrE-like derivation shows statistically robust sensitivity to p38-MAPKi (Fig. 5). Collectively, these ES-cell data support a role for p38-MAPK activity in facilitating PrE-like cell differentiation.

**Fig. 5:**
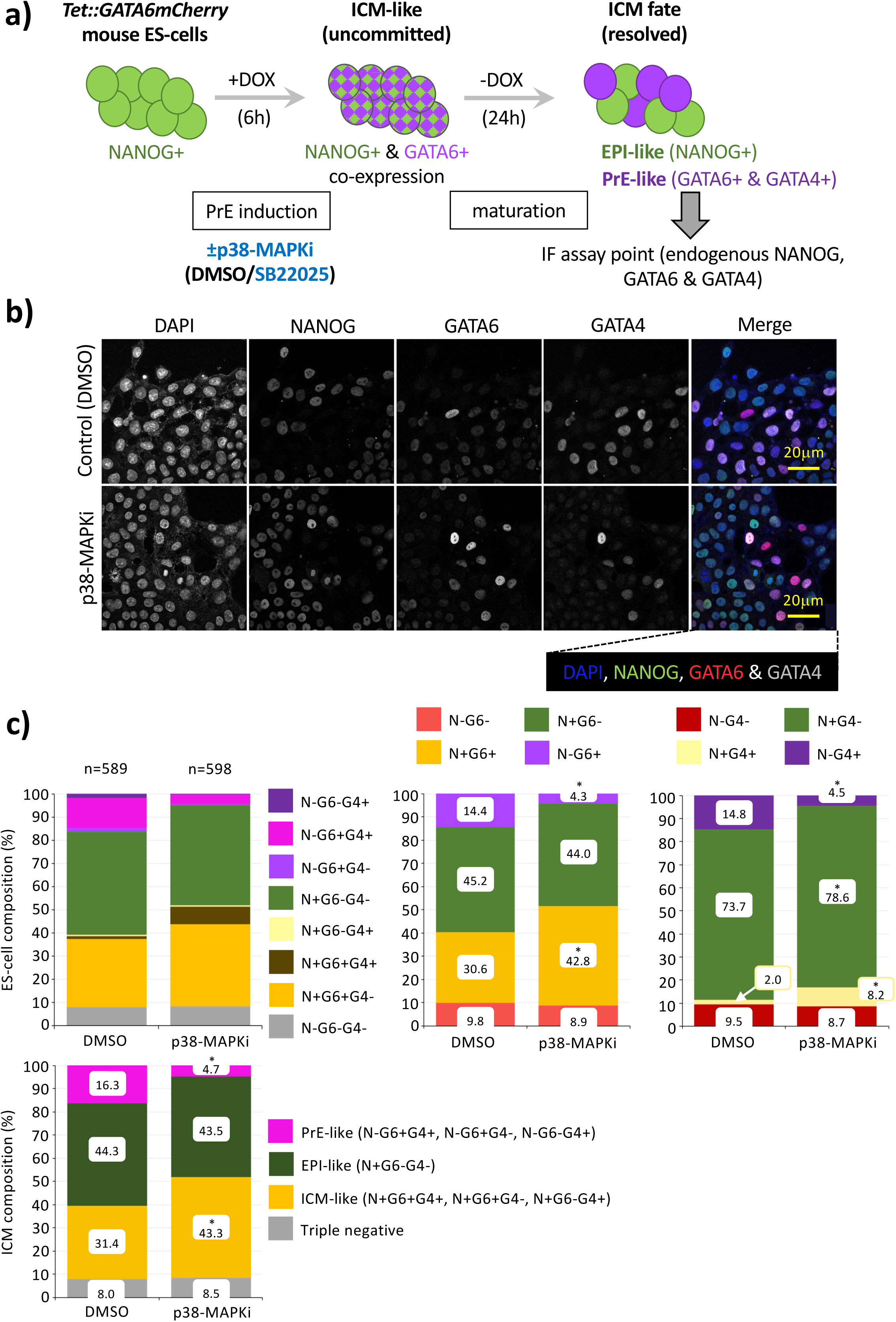
p38-MAPKi impairs PrE-like specification and differentiation in a 2D recombinant GATA6-inducible mouse ES-cell model: **a)** The mouse ES-cell line *Tet::Gata6mCherry* is induced by a 6-hour pulse of doxycycline treatment (+DOX 6h) to co-express endogenous NANOG and a recombinant GATA6-mCherry. After another 24 hours of culture in regular culture media (-DOX 24h), individual cells mature to express NANOG or endogenous GATA6/GATA4 in a mutually exclusive pattern (Schroter et al. 2015). ES-cell cultures were exposed to +DOX 6h induction plus p38-MAPKi (10μM SB220025) or vehicle control (DMSO) before transfer into regular culture -DOX 24h conditions for 24 hours, followed by fixation and IF-staining (NANOG, GATA6 & GATA4). **b)** Example ES-cell confocal micrographs related to the indicated +DOX 6h treatment conditions, IF stained for endogenous NANOG, GATA6 and GATA4 (grey-scale); plus pseudo-coloured merge image with DAPI counterstain. Scale bars - 20μm. **c)** Average cellular percentage composition of ICM cell lineage marker protein (co-)expression of indicated culture conditions, detailing; i. all possible cell lineage marker expression combinations (upper-left panel; +/-N = NANOG, +/-G6 = GATA6 & +/-G4 = GATA4), ii. compared NANOG & GATA6 (co-)expression status (upper-mid panel), iii. compared NANOG & GATA4 (co-)expression status (upper-right panel) and iv. classified as blastocyst PrE-like, EPI-like or ICM-like (uncommitted) cell lineages and triple-negative cells (lower left panel). Individual percentage ICM contributions and statistically significant differences are highlighted (Z-test; *p<0.05; total ES-cell n-numbers highlighted).

An additional feature of the *Tet::GATA6mCherry* ES-cell system is that after recombinant GATA6mCherry induction, cells can be driven to form 3D aggregates that, following 48 hours of culture, develop into ICM organoid structures with an outer PrE-like layer (expressing GATA6, SOX17, and GATA4) surrounding an inner population that expresses the EPI marker NANOG (Mathew et al. 2019). When we formed ICM organoids from induced *Tet::GATA6mCherry* ES cells cultured under control or p38-MAPKi conditions, we observed significant reductions in the percentage of PrE-like cells, but not to the same extent as observed in 2D cultures: N−G6+ declined from 21.1% (DMSO) to 18.9% (p38-MAPKi) and N−G4+ declined from 21.0% (DMSO) to 18.1% (p38-MAPKi). These reductions were accompanied by modest but significant increases in cells expressing neither NANOG nor GATA6/GATA4 (N−G6−: 12.8% to 16.2%; N−G4−: 12.9% to 17.0%) (Fig. S20, plus Fig. S21 with quantified protein expression data showing generally reduced marker levels in ICM organoids derived from p38-MAPKi-treated ES-cell induction, except for NANOG in N+G4+ cells). Taken together, these data suggest that while p38-MAPKi treatment impairs PrE differentiation from the derived uncommitted-like GATA6-mCherry–induced ES-cell population (Fig. 5 and Figs. S16 & S17), the prolonged 48-hour minus DOX culture required to form 3D aggregates provides a permissive window for partial compensatory cell-fate resolution, though complete restoration of PrE differentiation observed in controls is not achieved.

### Conclusions

We have reported that p38-MAPKi during mouse blastocyst maturation (E3.5–E4.5) robustly impairs PrE specification/differentiation and is associated with significantly reduced blastocyst cavity expansion ((Thamodaran and Bruce 2016; Bora et al. 2019; Bora et al. 2021b); recapitulated here in Figs. S1–S6. This cavity-expansion defect resembles data showing that experimentally induced cavity defects lead to decreased expression of ICM fate marker proteins (EPI–SOX2; PrE–GATA4; Ryan et al. 2019)). We sought to determine if p38-MAPKi–mediated PrE phenotypes could be reproduced by OA-induced cavity-volume impairments or whether they arise from autonomous ICM mechanisms. In agreement with the previous report (Ryan et al. 2019), OA-related cavity defects (Fig. 1) correlate with reduced EPI and PrE marker expression (Figs. S8 & S10); moreover, expanding these findings shows an associated impairment in specification from initially uncommitted ICM cells of overall EPI rather than PrE cell populations (Figs. 2–3 and Figs. S7, S9). These contrasting results indicate that while cavity expansion is impaired after p38-MAPKi, it is not the primary reason for the observed PrE deficit; instead, p38-MAPKi sensitivity involves ICM-autonomous mechanisms, as supported by observations in cultured IS-isolated ICMs (Fig. 4 and Fig. S15) and in both 2D and 3D mouse ES-cell/organoid ICM-related models (Fig. 5 and Figs. S16–S21). Thus, active p38-MAPK functions during early mouse blastocyst maturation to facilitate PrE specification/differentiation from uncommitted ICM progenitors as an intrinsic property of ICM cells, presumably responding to spatial–temporal cues that remain to be defined, culminating in the formation of the peri-implantation E4.5 blastocyst.

## DISCUSSION

It was previously reported that OA-mediated inhibition of ATP1 during mouse blastocyst maturation (E3.5–E4.0) impairs blastocyst cavity expansion and markedly reduces expression of both EPI (SOX2) and PrE (GATA4) markers (Ryan et al. 2019). We recapitulated OA-induced cavity defects (Fig. 1) and observed reduced ICM cell expression of an expanded set of EPI (NANOG, SOX2) and PrE (GATA6, GATA4, SOX17) markers by the late blastocyst stage (E4.5; Figs. S8 & S10). However, unlike Ryan and co-workers (Ryan et al. 2019), OA treatment from E3.5 to the mid-blastocyst (E4.0) stage did not uniformly reduce EPI and PrE marker proteins, at least in cells showing mutually exclusive EPI or PrE marker protein expression patterns (Figs. S8 & S10). This discrepancy may reflect our use of blastocysts *in vitro* cultured from the 2-cell (E1.5) stage versus freshly recovered E3.5 blastocysts (i.e. where the stated E4.0 stage may correspond more closely to a more advanced developmental stage akin to the E4.5 time-point in our data). Nonetheless, both datasets converge around our E4.5 time point, and together support mechanotransduction or trans-cavity signalling (possibly via luminal FGF4) as contributing to ICM lineage marker expression regulation (Ryan et al. 2019). Importantly, unlike the study of Ryan and colleagues, our expanded analyses, including the late E4.5 assay point and a broader panel of lineage markers (notably NANOG with GATA6), reveal that OA treatment primarily impairs EPI cell specification from uncommitted ICM, while PrE populations (GATA6/GATA4 without NANOG) are not similarly reduced (Fig. 2 and Fig. S7); a pattern opposite to the p38-MAPKi–induced PrE defects we described (Thamodaran and Bruce 2016; Bora et al. 2021b) and further confirmed here (again with an extended panel of lineage markers; Figs. S1 & S2). Our additional analysis of SOX2 and SOX17 expression reveals that OA treatment has specific effects regarding different EPI and PrE lineage markers, as by E4.5 there were no significant differences in the proportion or number of ICM cells expressing SOX2 and/or SOX17 (Fig. 3 and Fig. S9) compared to controls; in contrast to NANOG and GATA6 expression data (Figs. 2 and Fig. S7). However, exposing embryos to OA until the E4.0 stage did lead to a significant increase in the proportion and number of ICM cells existing in a previously unreported state, characterised by the co-expression of SOX2 and SOX17. While this conflicted state is also seen in control blastocysts, it occurs at a significantly lower frequency (Fig. 3 and Fig. S9). These results suggest that OA treatment enhances or prolongs this transient and conflicted cell fate state within the ICM. The increased frequency of SOX2 and SOX17 co-expression is associated with a reduction in the number of ICM cells expressing SOX2 alone, while the number of cells expressing only SOX17 remains unchanged (Fig. 3 and Fig. S9). This indicates that the enhanced conflicted cell fate state caused by OA treatment primarily affects the specification of epiblast (EPI) progenitors rather than primitive endoderm (PrE) progenitors. Specifically, in EPI progenitors, OA treatment disrupts the normal resolution process by which NANOG and SOX2 cease to be co-expressed with PrE markers (GATA6 and SOX17, respectively), with this disruption being more persistent for NANOG. This may reflect differences in the regulation or stability of GATA6 and SOX17, as by E4.5, quantitative analysis shows that in cells co-expressing EPI and PrE markers, GATA6 levels are similar to controls, but SOX17 levels are significantly reduced (Figs. S8 & S10). The OA-induced increase in NANOG and GATA6 co-expression likely indicates persistence of the uncommitted inner cell mass (ICM) state seen in E3.5 blastocysts (Chazaud et al. 2006), while the rise in SOX2 and SOX17 co-expression reflects enhanced induction of a conflicted, yet typically short-lived, cell fate state. Overall, our analysis of ICM cell fate in blastocysts with impaired cavity expansion shows that, both EPI and PrE marker protein expression levels per ICM cell are ultimately significantly reduced, consistent with previous findings (Ryan et al. 2019). Furthermore, that OA treatment specifically disrupts the appropriate specification of EPI cell populations, whether they arise from uncommitted cells (co-expressing NANOG and GATA6) or from fate-conflicted cells (co-expressing SOX2 and SOX17), while the specification and differentiation of PrE cells remains unaffected.

The EPI-specific ICM cell fate phenotypes associated with OA treatment are notably different from the PrE specification and differentiation defects previously observed after p38-MAPKi (Figs. S1, S2, S4 & S5; (Thamodaran and Bruce 2016; Bora et al. 2019; Bora et al. 2021b)), even though both inhibitors significantly reduce blastocyst cavity expansion (Fig. 1 and Figs. S3 & S6; Bora et al. 2021b). This indicates reduced cavity expansion after p38-MAPKi is unlikely to be the main cause of impaired PrE differentiation, and supports an ICM or ICM cell-autonomous mechanism, independent of TE-derived influence. Supporting this hypothesis, we found that treating isolated ICMs from early blastocysts with p38-MAPKi, thus eliminating any influence from cavity expansion, during the critical E3.5–E4.0 period of irreversible p38-MAPKi sensitivity (Thamodaran and Bruce 2016; Bora et al. 2021b), also leads to defective PrE specification and differentiation, while EPI specification remains unaffected (Fig. 4 and Fig. S15). Using established *in vitro* transgenic mouse ES-cell models of ICM cell fate resolution (Schroter et al. 2015; Mathew et al. 2019), we further demonstrate p38-MAPKi treatment effectively impairs the formation of a PrE-like lineage, while not affecting cells expressing EPI-like markers (Fig. 5 and Figs. S17 & S20). Notably, in the 2D culture model, six hours of p38-MAPKi exposure during the +DOX GATA6-mCherry induction period is sufficient to cause a significant reduction in the number of cells already initiating differentiation towards a PrE-like fate. This is accompanied by a significant increase in cells resembling the uncommitted early (E3.5) blastocyst ICM state (Fig. S17). This finding accords with our previous results on intact mouse blastocysts, which revealed a brief and irreversible window of PrE sensitivity to p38-MAPKi before the mid-blastocyst (E4.0) pattern of mutually exclusive NANOG and GATA6 expression emerges (Thamodaran and Bruce 2016; Bora et al. 2021b); strongly suggesting p38-MAPK activity is required early in the specification process. Furthermore, when the 2D ES-cell cultures were subsequently grown in regular - DOX media (after p38-MAPKi withdrawal) the same pattern persisted (Fig. 5). This indicates the initial six hours of p38-MAPKi treatment during induction is sufficient and irreversible in impairing PrE specification from uncommitted progenitors, up to at least 24 hours, reinforcing the conclusion that active p38-MAPK plays a crucial early role in PrE specification. Similarly, experiments testing the effect of p38-MAPKi on isolated ICMs demonstrated that early-stage inhibition, followed by a return to regular culture for 12 hours, was enough to block PrE specification and differentiation (Fig. 4). In the 3D ES-cell aggregation ICM organoid model, applying p38-MAPKi during the 6-hour +DOX induction period (before cell aggregation and the unavoidably necessary 48 hours of regular culture) significantly impaired the formation of PrE-like cells (identified by GATA6 and/or GATA4 expression without NANOG). However, this effect was less pronounced than in 2D cultures and did not result in a statistically significant increase in uncommitted cell types (co-expressing NANOG and GATA6); nevertheless, the proportion of EPI-like cells (NANOG-positive, GATA6-negative) remained consistently unaffected (Fig. S20). We speculate that the additional 24 hours of culture needed for ICM organoid formation, in the absence of p38-MAPKi, may allow for some compensatory regulation, which could explain the weaker effect compared to the 2D model. Attempts to analyse ICM lineage marker expression after 24 hours, to detect a possible increase in uncommitted PrE progenitors, were unsuccessful due to the instability of the ES-cell aggregates during fixation and staining. Notwithstanding, after 48 hours post-DOX induction with p38-MAPKi, there was a significant reduction in PrE-like cell formation and lower average per-cell expression of GATA6 and GATA4 (Fig. S21). In intact blastocyst cultures, p38-MAPKi treatment after E3.75 or E4.0 (until E4.5) does not impair PrE specification, whereas similar inhibition of MEK1/2 (using PD0325901, which blocks FGF4 signalling promoting PrE differentiation (Yamanaka et al. 2010)) robustly blocks PrE formation (Thamodaran and Bruce 2016). Collectively, these findings indicate active p38-MAPK plays an important and temporally early role in blastocyst development, prior to ICM lineage specification, that is distinct from the later FGF4/MEK1/2-dependent mechanisms; ultimately leading to irreversible ICM cell fate decisions by the late blastocyst (E4.5) stage (Nichols et al. 2009; Yamanaka et al. 2010; Frankenberg et al. 2011; Kang et al. 2013; Thamodaran and Bruce 2016).

We previously found that p38-MAPKi-mediated PrE phenotypes are associated with impaired general protein synthesis, reduced ribosomal protein expression, decreased polysome formation, and disrupted rRNA precursor processing. Moreover, p38-MAPKi-induced PrE defects could be partially rescued by pharmacologically activating the mTOR pathway (a key regulator of metabolism and protein synthesis (Fonseca et al. 2014)), without influencing impaired blastocyst cavity expansion (Bora et al. 2021b). This partial rescue suggested that p38-MAPK enables PrE specification and differentiation through mechanisms intrinsic to ICM cells, independent of cavity expansion, as supported by data herein. We propose that early blastocyst p38-MAPK signalling may ensure germane PrE-specification from uncommitted progenitors via ICM autonomous regulation of protein translation/synthesis; thereby priming such cells for differentiation. We recently reported that inhibition of mTORC1 during the 8- to 16-cell transition impairs the spatial allocation of daughter blastomeres to the emerging population of primary ICM founder cells; via a mechanism potentiating the 7-methyl-guanosine cap (m7G-cap) binding complex (EIF4F), which is essential for efficient translation of mRNA subclasses containing 5’UTR terminal oligopyrimidine (TOP) motifs. A phenotype replicated by similar p38-MAPKi treatment (Gahurova et al. 2023). p38-MAPKi effects on blastocyst PrE specification may also be related to similar impairments in selective mRNA translation, which could be readily investigated using embryo optimised SSP-ribosomal profiling (Del Llano et al. 2020; Masek et al. 2020; Bora et al. 2021b; Iyyappan et al. 2023) or Ribo-ITP-RNAseq (Ozadam et al. 2023) to identify differential mRNA ribosome-association under ±p38-MAPKi conditions. However, p38-MAPKi-induced PrE differentiation defects are also characterised by increased numbers of uncommitted ICM cells co-expressing both GATA6 and NANOG (Figs. S1 & S2; Thamodaran and Bruce 2016; Bora et al. 2021b), suggesting that p38-MAPK activity is also required to clear putative PrE progenitors of pluripotency-associated NANOG expression, potentially through mechanisms related to protein stability. Indeed, mTORC1 inhibition from E3.5 also increases the number of cells co-expressing NANOG and GATA6, although this is associated with general developmental diapause (Bulut-Karslioglu et al. 2016; Bora et al. 2021b). This point is further supported by the observation of significant NANOG and GATA4 co-expression in p38-MAPKi-treated blastocysts exposed to concurrent mTOR activation (Bora et al. 2021b), whereby PrE specification is partially rescued but NANOG expression endures. Thus, the ICM-autonomous role of p38-MAPK in promoting PrE specification and differentiation is likely to be multi-factorial.

In summary, our findings show that OA-induced blastocyst cavity expansion leads to reduced marker protein expression in both ICM lineages, but specifically impairs the specification of uncommitted ICM progenitors towards the EPI lineage. This is the opposite of the effect observed after p38-MAPKi during the same developmental period, despite similar cavity expansion defects. Further, p38-MAPKi treatment of isolated ICMs and mouse ES-cell models consistently impairs PrE cell differentiation, while EPI lineages remain largely unaffected. Together, these results highlight an ICM-autonomous role for active p38-MAPK in priming PrE specification and differentiation during early blastocyst maturation, possibly via specific regulation of mRNA translation, and distinct from mechanosensory mechanisms linked to cavity expansion.

## MATERIALS & METHODS

### Superovulation and embryo isolation

Animal work was conducted in accordance with Act No 246/1992 Coll., on the protection of cruelty against animals under the supervision of the Central Commission for Animal Welfare, approval ID 51/2015 (Czech Republic). Experimental embryos were collected as previously described (Mihajlovic et al. 2015). Briefly, F1 generations of 8-week female hybrid mice (C57BL6 female and CBA/W strain crosses) were intraperitoneally injected with 7.5IU of PMSG (pregnant mare serum gonadotrophin; Merck) and reinjected after 48 hours with 7.5IU hCG (human chorionic gonadotrophic hormone; Merck), before overnight mating with F1 stud males. 4 hours before dissection, culture dishes containing twenty 10μl drops of KSOM+AA medium (Embryo-Max; Millipore), covered with mineral oil (Irvine Scientific) and equilibrated at 37°C in a 5% CO_2_ atmosphere were prepared. For inhibitor treatments on intact blastocysts, recovered 2-cell stage embryos (E1.5; 45-47 hours post-hCG) were washed through 20μL drops of prewarmed (37°C) M2 media (EMD Millipore Corp. – 3816047) containing 4mg/ml BSA (bovine serum albumin - Merck) and then transferred through a series of prewarmed KSOM+AA culture drops and *in vitro* cultured to the desired stage; including exposure to control or specific inhibitor treatments (see below). For immuno-surgery (IS) experiments, 8-cell (E2.5) embryos were recovered and similarly cultured in KSOM+AA to the desired early-blastocyst (E3.5) stage for ICM isolation (91 or 96 hours post-hCG injection – see below) and further *in vitro* cultured under control or inhibitor treatment conditions.

### Immuno-surgical (IS) ICM isolation

A previously described protocol was adopted (Wigger et al. 2017). Briefly, early stage (E3.5) blastocysts (at a stage equivalent to 91 or 96 hours post-hCG injection), obtained from *in vitro* cultured embryos recovered at the 8-cell (E2.5) stage, were subject to *zona pellucidae* removal in prewarmed (37°C) acid Tyrode’s Solution drops (Sigma-Aldrich. cat. #T1788) diluted in M2. Blastocysts were transferred to M2 media drops for 15 minutes, incubated for 40 minutes in anti-mouse serum antibody (diluted 1:2 in M2 media: Merck – M5774) and then guinea pig complement (1:4 in prewarmed M2 media) for 30 minutes; initiating complement-mediated outer TE cell lysis. ICMs were isolated by repeated pipetting in prewarmed M2 media drops and subject to immediate fixation and IF staining (see below) or cultured in KSOM+AA to the E4.5 equivalent developmental stage.

### Embryo/ IS isolated ICM inhibitor treatments

Ouabain-treatment; Early- (E3.5) staged blastocysts (with cavities approximating half the embryonic volume) were transferred into pre-equilibrated KSOM+AA culture drops containing either 500μM Ouabain (Merck: O3125) or equivalent volumes of vehicle dimethylsulfoxide (DMSO, Merck: D4540) solvent control and cultured to the mid- (E4.0) or late- (E4.5) blastocyst stages, fixed and, as necessary, IF stained (see below). p38-MAPKi using SB220225 (Calbiochem: #559396); as for OA-treatment utilising 20μM SB220225 (Merck: 559396), unless otherwise stated, or equivalent volumes of DMSO control. IS isolated ICMs p38-MAPKi treatments; isolated ICMs were cultured (5, 10 or 20μM SB220225 or equivalent volumes of DMSO control) to the E4.0 equivalent stage, before transfer to non-supplemented KSOM+AA and culture to E4.5, prior to fixation and IF staining.

### Blastocyst/IS isolated ICM fixation and immuno-fluorescent (IF) staining

Blastocyst/ICM IF staining protocols are described in (Mihajlovic et al. 2015). Briefly, blastocyst *zona pellucidae* were removed in prewarmed (37°C) drops of Acid Tyrode’s solution diluted in M2. Blastocysts/ICMs were fixed, on 1.5% agar-coated culture dishes, in 20μl drops of a 4% para-formaldehyde solution (PFA; Santa Cruz Biotechnology), overlaid with mineral oil, for 20 minutes at 37°C. All subsequent steps were performed at room temperature unless stated. In 96-well micro-titre plates, blastocysts/ICMs were washed through three 70μl drops of PBST (phosphate-buffered saline with 0.15% Tween 20 - Merck), placed in 50μl of 0.5% Triton-X100 (Merck) permeabilisation solution (diluted in PBS) for 20 minutes, washed through three 70μl drops of PBST and transferred to 50μl drops of blocking 3% BSA (Merck) in PBST for 30 minutes. Desired primary antibody dilutions (see below) were prepared in 3% BSA PBST solutions (in 5μL volumes overlaid with mineral oil) for overnight incubation at 4°C. After three 70μL drop washes of PBST, embryos/ICMs were subject to a secondary 3% BSA block (1 hour) and transferred into 5μl 3% BSA drops containing appropriate dilutions of fluorescently-conjugated secondary antibody (see below) and incubated in the dark at 4°C for 3 hours. After three further 70μl PBS-T washing steps, embryos/ICMs were DNA counter-stained using Vectashield mounting media containing DAPI (Vector). Primary antibodies (and dilutions): a) raised in rabbit: anti-GATA4 (sc-9035, Santa Cruz Biotechnology - 1:200), b) raised in mouse: anti-SOX2 (sc-365823, Santa Cruz Biotechnology - 1:200), CDX2 (MU392A-UC; BioGenex – 1:200), c) raised in goat: anti-SOX17 (AF1924, R&D Systems – 1:200), anti-GATA6 (AF1700, R&D Systems – 1:200) and d) raised in rat: anti-NANOG (14-5761, Affymetrix/eBiosciences – 1:200). Secondary antibodies (and dilutions): i. donkey anti-rabbit-Alexa-Fluor^647^ (ab150075, Abcam 1:500), ii. donkey anti-mouse-Alexa-Fluor^488^ (Abcam # ab150107, ThermoFisher - 1:500), iii. donkey anti-goat-Alexa-Fluor^555^ (A214232, ThermoFisher - 1:500) and iv. donkey anti-rat-Alexa-Fluor^488^ (A21208, ThermoFisher – 1:500).

### Blastocyst/IS isolated ICM confocal microscopy & image analysis *(inc. statistics)*

IF stained blastocysts/ICMs were imaged, using standard protocols, by fluorescent confocal microscopy as previously described (Mihajlovic et al. 2015; Mihajlovic and Bruce 2016; Thamodaran and Bruce 2016; Bora et al. 2019; Bora et al. 2021a; Bora et al. 2021b; Gahurova et al. 2023); ensuring identical acquisition settings for all comparative control versus experimental groups (also simultaneously IF stained). Briefly, blastocyst/ICM samples were imaged on glass-bottomed 35mm culture dishes (NC9341562, MatTek), in 20μl drops of Vectashield mounting media overlaid with mineral oil, on an inverted Olympus FV10i confocal microscope. Per blastocyst/ICM, a full and non-overlapping z-series of confocal sections was acquired and micrograph images visualised (and prepared for figure generation) using the Olympus Fluoview ver.1.7, FIJI (Schindelin et al. 2012) or Imaris software. Per blastocyst/ICM, individual cell numbers were counted (based on DAPI-stained nuclei) and further subcategorised as, outer or inner cells, and then based on their composite expression of detectable IF-stained lineage markers (see tabulated summaries – supplementary Excel data/stats tables). Absolute mean cell numbers, per blastocyst/ICM, for each lineage marker expressing category were calculated as average percentage contribution to the overall ICM populations in each control and experimental condition derived (an infrequent minority of mitotic ICM cells were excluded; such absolute numbers (plus calculated mean standard deviations) and percentage contributions were charted using GraphPad Prism 8 or Microsoft Excel, respectively. Statistical significance of control versus experimental conditions were determined; i. for normally or non-normally distributed absolute cell number mean data sets (determined with D’Agostino & Pearson and Anderson-Darling tests), using unpaired student t-tests or Mann-Whitney tests, respectively, and ii. for average percentage ICM contribution using appropriate Z-tests or Mann-Whitney tests (or in the case of IS ICM-related data, using unpaired student t-tests) employing a significance threshold of p<0.05 (*); see supplementary Excel data/stats tables. Individual cell lineage nuclear marker protein expression levels, per blastocyst/IS isolated cultured ICM (from four randomly selected, blastocysts/ICMs in each quartile with respect to total blastocyst cell number) were calculated, as previously described (Potapova et al. 2011; McCloy et al. 2014) and employed (Bora et al. 2021b; Gahurova et al. 2023), as corrected total cell fluorescence (CTCF) in FIJI (Schindelin et al. 2012); from minimal composite confocal micrograph z-sections (encompassing the entire nucleus). Individual measurements were set using the FIJI command Analyze>Set Measurements, and the following options selected: Area, Mean grey value and Integrated Density. Using the Polygon selection tool, an area encompassing individual cell nuclei were demarcated, and measurements recorded. The selected area was translocated to encompass an area excluding the embryo for normalising background measurements. From repetitive tabulated data, the corrected total cell fluorescence (CTCF), in arbitrary units, was determined for each nucleus as follows: CTCF = integrated density – (area of selected cell × mean fluorescence of background readings). Calculated CTCFs were plotted as scatterplots, indicating means and standard deviations (GraphPad Prism 8) and mean CTCF differences statistically tested via Mann-Whitney tests or unpaired t-tests after confirmation of normally distributed data (D’Agostino & Pearson and Anderson-Darling tests); significance threshold p<0.05 (*)(*); see supplementary Excel data/stats tables.

### Blastocyst cavity volume calculations

For Ouabain and SB220225 treated (plus corresponding DMSO controls) fixed blastocyst groups, the pico-litre (pL) volume of derived cavities was determined in FIJI (ImageJ (Schindelin et al. 2012)) by measuring the outer circumference of the cavity in the centrally located widest Z-stack (with the following settings: Analyze>Set Measurements; and selecting the Perimeter option; using the Polygon selection tool, to trace and measure the inner cavity circumference). The radii of measured circumferences were deduced and used to calculate an approximate equatorial sectional area from which a pL value could be mathematically determined. Standard deviations were calculated, and observed differences in the experimental/control cohorts were statistically tested (after assessing the normal or non-normal distribution of the data with D’Agostino & Pearson and Anderson-Darling tests) by using unpaired student t-tests or Mann-Whitney tests, respectively (*p<0.05) (*); see supplementary Excel data/stats tables.

### 2D mouse ES-cell (ESC) & 3D ICM organoid model cultures and p38-MAPKi

Mouse ES-cells (*Tet::*GATA6mCherry) were maintained on 0.1% gelatine-coated tissue culture dishes in GMEM-based medium supplemented with 10% FBS, sodium pyruvate, 50µM β-mercaptoethanol, glutamax, non-essential amino acids and LIF; and incubated at 37°C and 5% CO_2_. To model ICM specification into EPI-like and PrE-like cells, 2D cultures of *Tet::*GATA6mCherry mouse ES-cells (Schroter et al. 2015), or derived of 3D ICM organoids (Mathew et al. 2019), were employed as described. For the 2D model, differentiation was induced by adding 500 ng/ml doxycycline (DOX) to the culture medium for 6 hours, followed by further culture in serum + LIF for additional 24 hours (in the absence of DOX). 3D ICM organoids were generated by a similar 6-hour induction in DOX-containing media, after which 50 ES-cells were seeded in coated (1% low-melt agarose in PBS) concave wells of 96-well plates in media containing serum + LIF (briefly pulsed centrifgued) and cultured for 48 hours (until aggregate ICM organoids formed). p38-MAPK inhibitor SB220025 at 5, 10 or 20μM (or equivalent volumes of DMSO control) was added during the + DOX induction phase for 6 hours (+doxycycline). IF staining was performed as previously described (Kalmar et al. 2009). Briefly, adhered cells or ICM organoids were rinsed in BBS-CaCl_2_ prior to fixation in 4% PFA for 15 minutes at room temperature and three washes in BBT-BSA (0.1% Triton X-100, 0.5% BSA) permeabilisation/blocking solution for 15 minutes. Cells were incubated in primary antibody overnight at 4°C in a humidity chamber. Primary antibodies included: NANOG (14-5761-80, eBIOSCIENCE - 1:200), GATA6 (AF1700, R&D Systems - 1:200), GATA4 (sc-9053, Santa Cruz Biotechnology - 1:200). Cells were then washed three times with BBT-BSA for 15 minutes before incubation in appropriate secondary antibody for 1-3 hours in the dark; i. donkey anti-rat Alexa-Fluor^488^ (A-21208, ThermoFisher - 1:1000), ii. donkey anti-goat Alexa-Fluor^568^ (A-11057, ThermoFisher – 1:1000), iii. donkey anti-rabbit Alexa-Fluor^647^ (A-31573, ThermoFisher – 1:1000). Nuclei were visualized via DAPI counter-staining (D1306, ThermoFisher - 1:1000 in PBS). Cells were then washed three times (10 minutes) in BBS-CaCl_2_ and mounted using prepared mounting medium (HMM, 4% N-propyl-galate, 80% glycerol).

### 2D mouse ES cell (mESCs) & 3D ICM organoid model microscopy and image analysis

Cells were imaged using an inverted Zeiss LSM Zeiss LSM700 and a Plan-Apochromat 40x/1.3 Oil DIC (UV) VIS-IR M27 objective. Images were acquired using 512 x 512 pixels (159.73μm x 159.73μm) resolution of specimens that were simultaneously IF stained under identical conditions and imaged under identical conditions regarding laser intensity, gain and pinhole aperture during a single confocal session. Zeiss AIM software (Carl Zeiss Microsystems) and FIJI (Schindelin et al. 2012) were used for image acquisition and visualization, respectively. Cell images were processed and analysed as previously described (Fischer et al. 2020). Briefly, a Matlab-based software called Modular Interactive Nuclear Segmentation (MINS) was used to segment ES-cell confocal images based on the DAPI nuclear stain (Lou et al. 2014). MINS detected and segmented each nucleus, providing a fluorescence intensity output of each channel of each individual cell. Composite analyses were performed, using the software Paleontological Statistics (PAST; https://palaeo-electronica.org/2001_1/past/issue1_01.htm), of NANOG, GATA6 and GATA4 fluorescence levels to permit the calculation of threshold levels by which individual cells were classified as positive or negative for each protein marker; individual cells were classified as either; i. triple negative (TN, negative expression for NANOG, GATA6 or GATA4; N-G6-G4-), ii. EPI progenitors (positive for NANOG and negative for GATA6 and/or GATA4; N+G6-, N+G4- or N+G6-G4-), iii. PrE progenitors (negative for NANOG and positive for GATA6 or GATA4; N-G6+, N-G4+ or N-G6+G4+) or iv. uncommitted cells (positive for both, NANOG and GATA6 or NANOG and GATA4; N+G6+ or N+G4+).

### 2D mouse ES cell (mESCs) & 3D ICM organoid model statistics

ANOVA with Tukey’s multiple comparison tests were used for comparisons between groups and Z-tests for frequencies, followed by the Bonferroni correction for multiple comparisons. Values of *p*<0.05 (*) were considered statistically significant. Statistical tests were performed using GraphPad Prism version 8.0.0 for Windows, GraphPad Software, San Diego, California USA (www.graphpad.com).

## Supporting information

Supplementary data/stats Excel tables

## ACKNOWLEDGEMENTS

We acknowledge the Institute of Parasitology (Biology Centre of the Czech Academy of Sciences, in České Budějovice) for housing mice, Marta Gajewska (Institute of Oncology, Warsaw, Poland) and Anna Piliszek (Institute of Genetics and Animal Breeding, Polish Academy of Sciences, Jastrzębiec, Poland) for founder CBA/W mice and other laboratory members for valuable inputs and discussions. Imaging was carried out at the SIMACE (Advanced Confocal and Electron Microscopy Research Service) at the IUIBS.

## SUPPLEMENTARY FIGURE LEGENDS

**Fig. S1:**
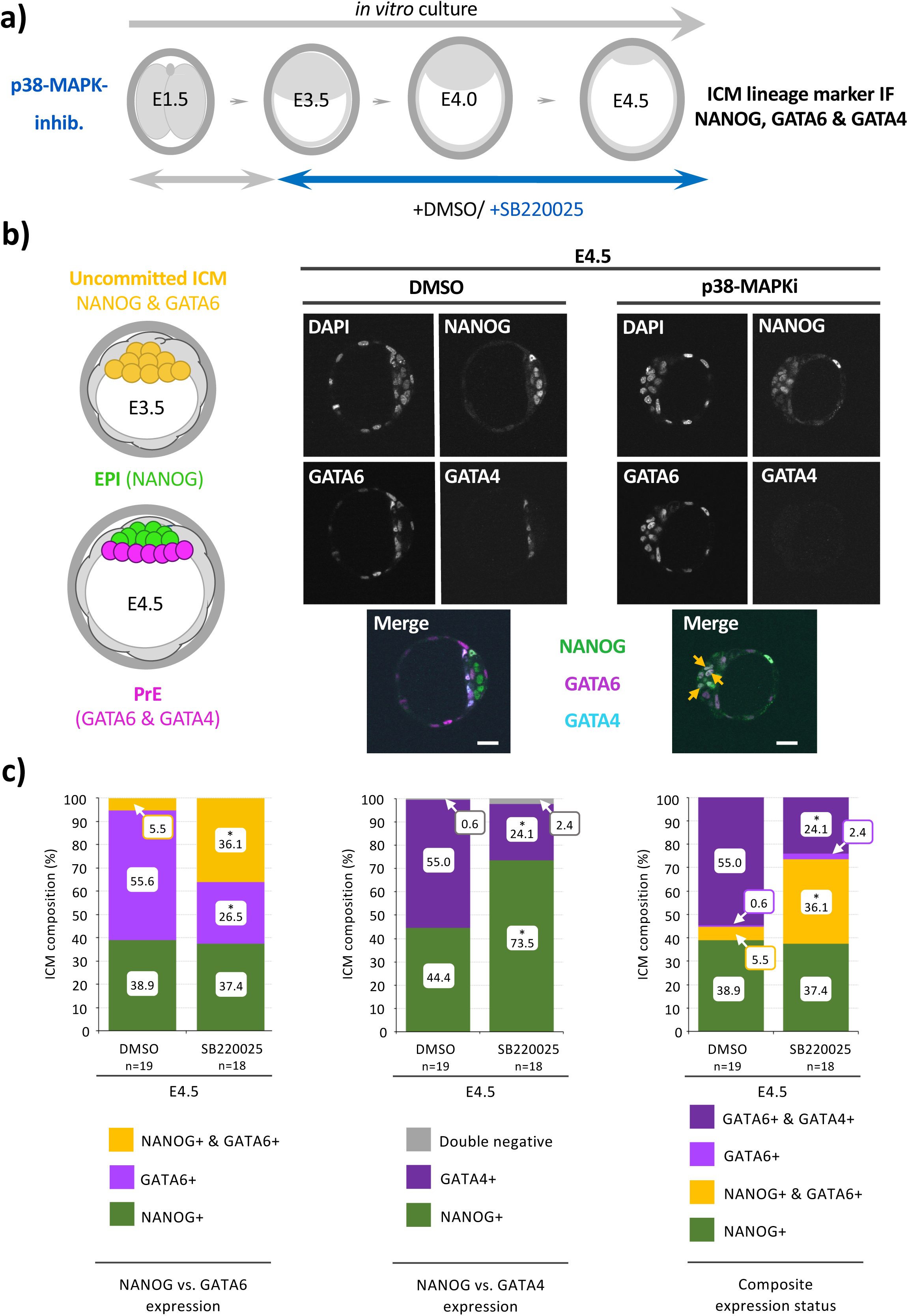
p38-MAPK inhibition (p38-MAPKi - SB220025) treatment during blastocyst embryo maturation (E3.5-E4.5) impairs PrE specification and differentiation from uncommitted ICM progenitors (IF: NANOG, GATA6 & GATA4 – exemplar confocal micrographs and average percentage population data): **a)** Experimental scheme of p38-MAPKi (SB220025 – 20μM) during blastocyst maturation period (E3.5-E4.5). **b)** Left -Schematic detailing typical expression of ICM lineage marker proteins in the uncommitted early-blastocyst (E3.5) stage (NANOG & GATA6 co-expression - yellow) and at the late-blastocyst (E4.5) stage (EPI – NANOG alone – green, PrE – GATA6 & GATA4 in the absence of NANOG - magenta). Right - Exemplar confocal single z-section micrographs of control (DMSO) or p38-MAPKi treated blastocysts, IF assayed for combined NANOG, GATA6 & GATA4 expression at E4.5 (scale bar - 20μm). **c)** Average cellular percentage ICM composition of control (DMSO) or p38-MAPKi treated blastocysts IF assayed for cell lineage marker expression; delineated as i. uncommitted (co-expressing NANOG & GATA6), PrE (expressing GATA6 without NANOG) and EPI (expressing NANOG without GATA6) cells (left panel), ii. PrE (expressing GATA4 without NANOG) and EPI (expressing NANOG without GATA4) cells (central panel), or iii. uncommitted (co-expressing NANOG & GATA6), PrE (expressing GATA6 alone or GATA6 plus GATA4, each without NANOG) and EPI (expressing NANOG without GATA6 or GATA4) cells (right panel). Statistically significant differences highlighted (Z-test or Mann-Whitney as appropriate; *p<0.05, see supplementary data/stats Excel tables; embryo n-numbers highlighted). *See Fig. S2 for average cell number population data/ per assayed embryo.* Note, p38-MAPKi results in unaffected ICM specification of EPI (denoted by NANOG expression in the absence of either GATA6 or GATA4), impaired formation of PrE (classified by GATA6 or GATA4 expression in the absence of NANOG) and increased incidence of uncommitted PrE progenitor cells (co-expressing NANOG and GATA6 – see orange arrows in panel b), as previously described (Thamodaran and Bruce 2016; Bora et al. 2019; Bora et al. 2021b).

**Fig. S2:**
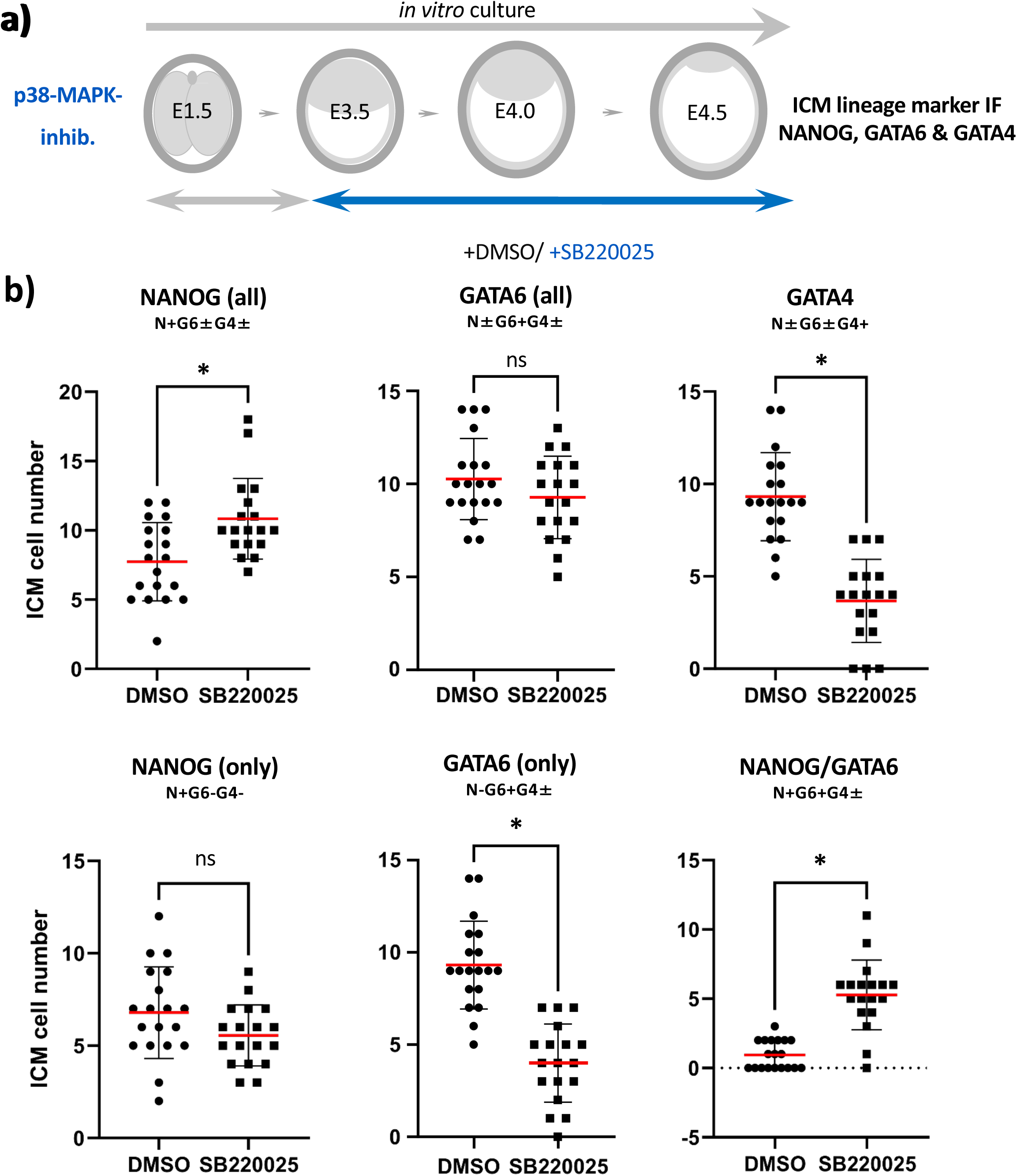
p38-MAPK inhibition (p38-MAPKi - SB220025) treatment during blastocyst maturation (E3.5-E4.5) impairs PrE specification and differentiation from uncommitted ICM progenitors (IF: NANOG, GATA6 & GATA4 – average cell number population data): **a)** Experimental scheme of p38-MAPKi (SB220025 – 20μM) during blastocyst maturation period (E3.5-E4.5). **b)** Average numbers of total NANOG, total GATA6, total GATA4, NANOG only (without GATA6 or GATA4), GATA6 only (without NANOG, ±GATA4) and NANOG & GATA6 co-expressing (±GATA4) ICM cells, per embryo, assayed at the late-blastocyst (E4.5) stage under control (DMSO) or p38-MAPKi treatment conditions; arithmetic means (red bars), non-statistical differences (denoted by ‘ns’) and statistically significant differences indicated by significance markers (for the non-normally distributed data a Mann-Whitney test was employed; *p<0.05, see supplementary data/stats Excel tables).

**Fig. S3:**
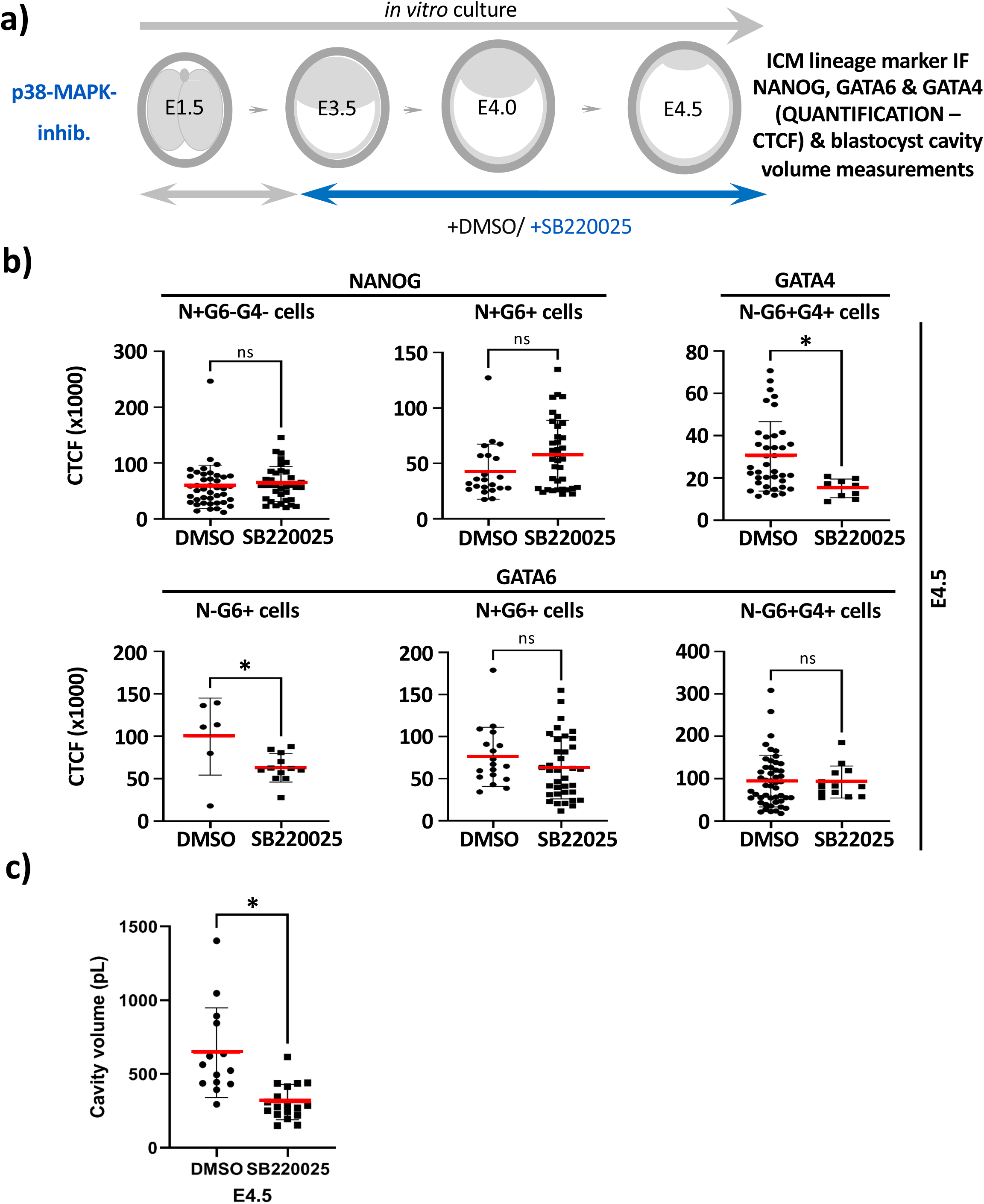
Quantification of average per blastocyst ICM cell expression levels of lineage protein marker expression, and blastocyst cavity volume, after p38-MAPKi treatment (E3.5-E4.0/E4.5; NANOG, GATA6 & GATA4): **a)** Experimental scheme of p38-MAPKi (SB220025 – 20μM) treatment and assay points during blastocyst maturation period (E3.5-E4.5). **b)** Average quantified ICM cell lineage marker expression (NANOG, GATA6 & GATA4; CTCF) in blastocysts cultured from E3.5-E4.5 under control (DMSO) and p38-MAPKi-treated conditions, according to indicated co-expression status; arithmetic means (red bars), standard deviations and statistically significant differences are highlighted (for normal or non-normally distributed data unpaired T-test or Mann-Whitney tests were employed, respectively; *p<0.05, see supplementary data/stats Excel tables). **c)** Measured blastocyst cavity volumes in i blastocysts cultured from E3.5-E4.5 under control (DMSO) and p38-MAPKi-treated conditions (pL); arithmetic means (red bars), standard deviations and statistically significant differences are highlighted (for the non-normally distributed data, a Mann-Whitney test was employed. *p<0.05; see supplementary data/stats Excel tables).

**Fig. S4:**
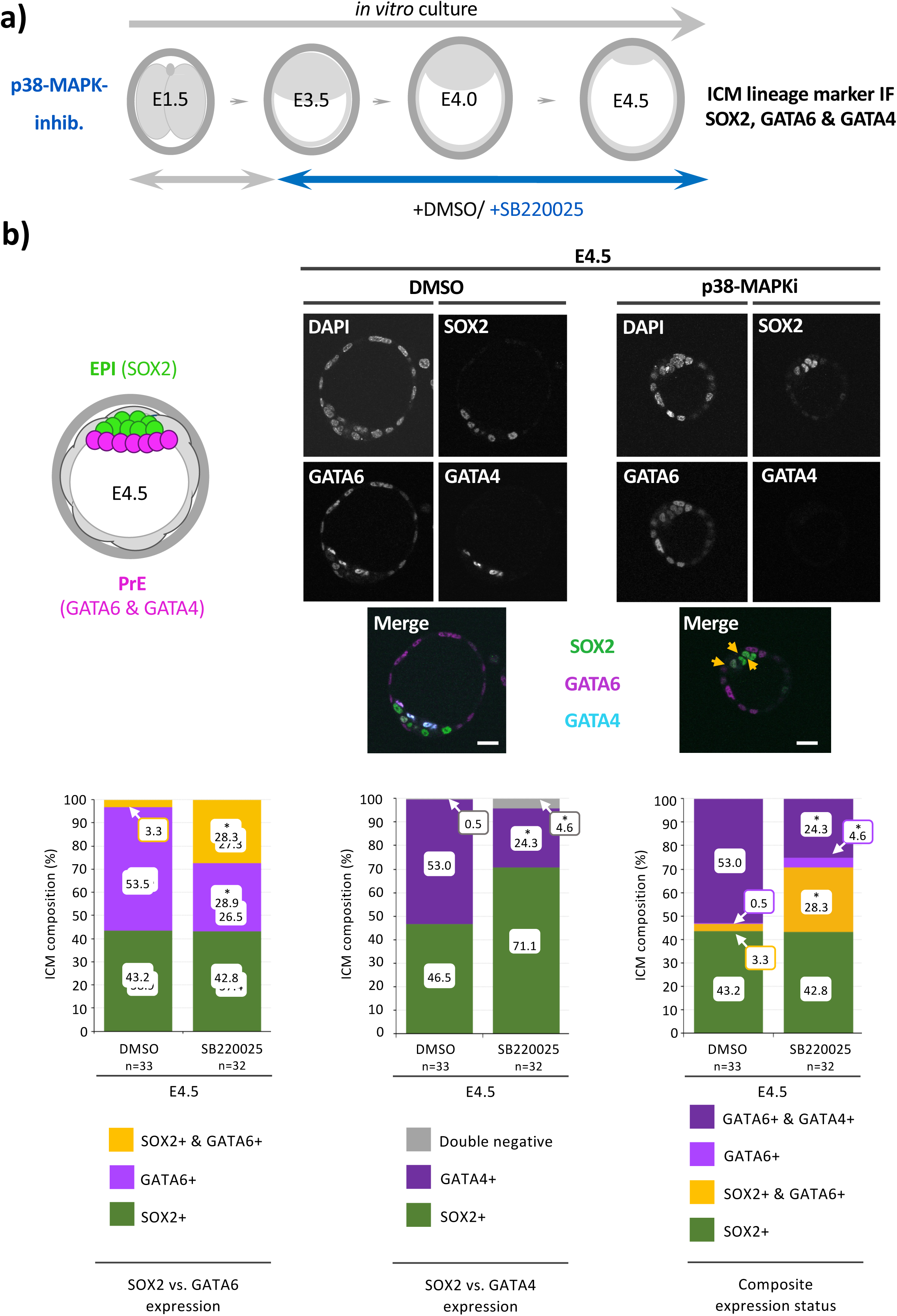
p38-MAPK inhibition (p38-MAPKi - SB220025) treatment during blastocyst embryo maturation (E3.5-E4.5) impairs PrE specification and differentiation from uncommitted ICM progenitors (IF: SOX2, GATA6 & GATA4 – exemplar confocal micrographs and average percentage population data): **a)** Experimental scheme of p38-MAPKi (SB220025 – 20μM) during blastocyst maturation period (E3.5-E4.5). **b)** Left -Schematic detailing typical expression of ICM lineage marker proteins at the late-blastocyst (E4.5) stage (EPI – SOX2 alone – green, PrE – GATA6 & GATA4 in the absence of SOX2 - magenta). Right - Exemplar confocal single z-section micrographs of control (DMSO) or p38-MAPKi treated blastocysts, IF assayed for combined SOX, GATA6 & GATA4 expression at E4.5 (scale bar - 20μm). **c)** Average cellular percentage ICM composition of control (DMSO) or p38-MAPKi treated blastocysts IF assayed for cell lineage marker expression; delineated as i. uncommitted/conflicted (co-expressing SOX2 & GATA6), PrE (expressing GATA6 without SOX2) and EPI (expressing SOX2 without GATA6) cells (left panel), ii. PrE (expressing GATA4 without SOX2) and EPI (expressing SOX2 without GATA4) cells (central panel), or iii. uncommitted/conflicted (co-expressing SOX2 & GATA6), PrE (expressing GATA6 alone or GATA6 plus GATA4, each without SOX2) and EPI (expressing SOX2 without GATA6 or GATA4) cells (right panel). Statistically significant differences highlighted (Z-test or Mann-Whitney as appropriate; *p<0.05, see supplementary data/stats Excel tables; embryo n-numbers highlighted). *See Fig. S5 for average cell number population data/ per assayed embryo.* Note, p38-MAPKi results in unaffected ICM specification of EPI (denoted by SOX2 expression in the absence of either GATA6 or GATA4), impaired formation of PrE (classified by GATA6 or GATA4 expression in the absence of SOX2) and increased incidence of uncommitted/conflicted PrE progenitor cells (co-expressing SOX2 and GATA6 – see orange arrows in panel b).

**Fig. S5:**
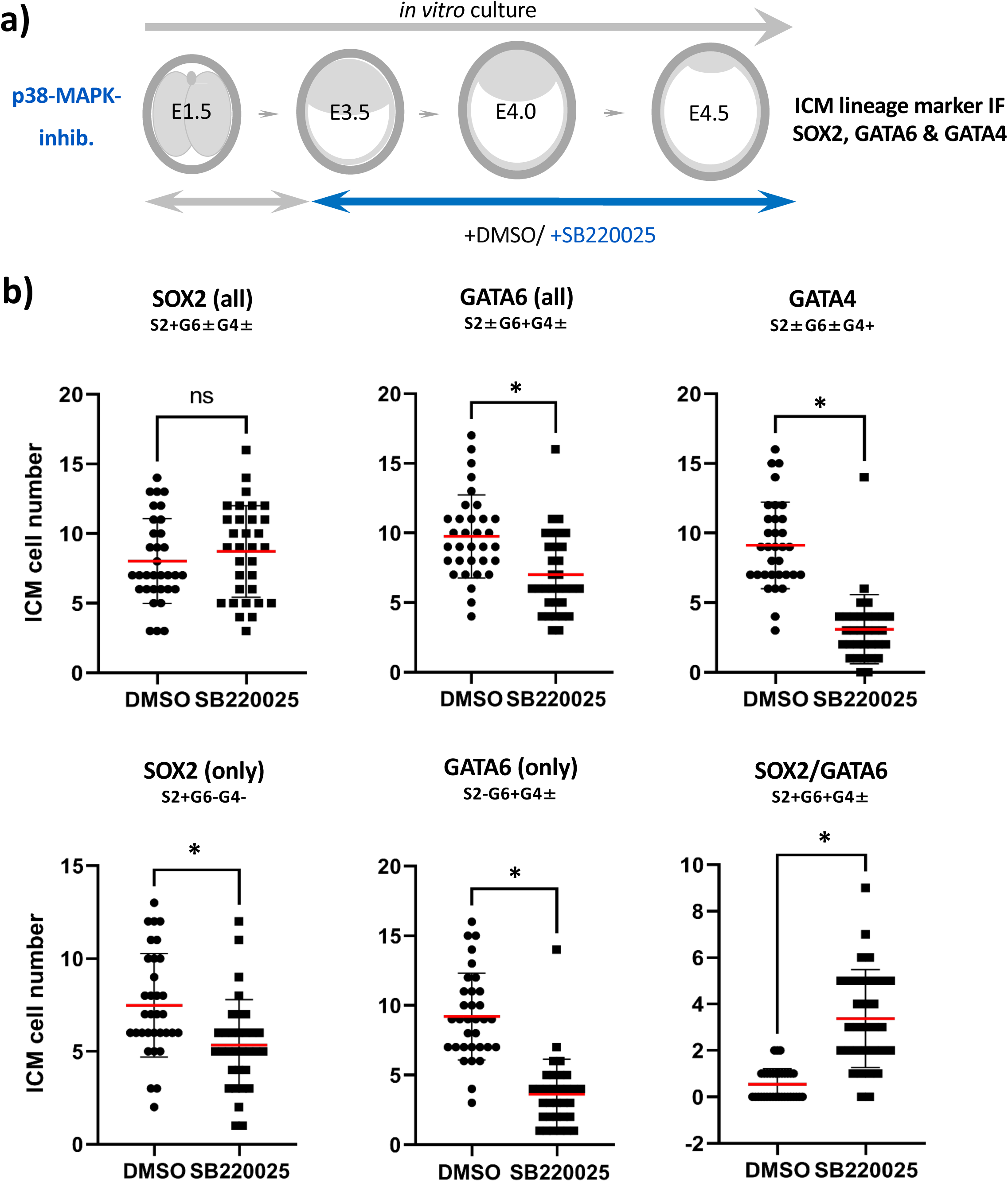
p38-MAPK inhibition (p38-MAPKi - SB220025, 20μM) treatment during blastocyst maturation (E3.5-E4.5) impairs PrE specification and differentiation from uncommitted ICM progenitors (IF: SOX2, GATA6 & GATA4 – average cell number population data): **a)** Experimental scheme of p38-MAPKi (SB220025 – 20μM) during blastocyst maturation period (E3.5-E4.5). **b)** Average numbers of total SOX2, total GATA6, total GATA4, SOX2 only (without GATA6 or GATA4), GATA6 only (without SOX2, ±GATA4) and SOX2 & GATA6 co-expressing (±GATA4) ICM cells, per embryo, assayed at the late-blastocyst (E4.5) stage under control (DMSO) or p38-MAPKi treatment conditions; arithmetic means (red bars), non-statistical differences (denoted by ‘ns’) and statistically significant differences indicated by significance markers (for the non-normally distributed data a Mann-Whitney test was employed; *p<0.05, see supplementary data/stats Excel tables).

**Fig. S6:**
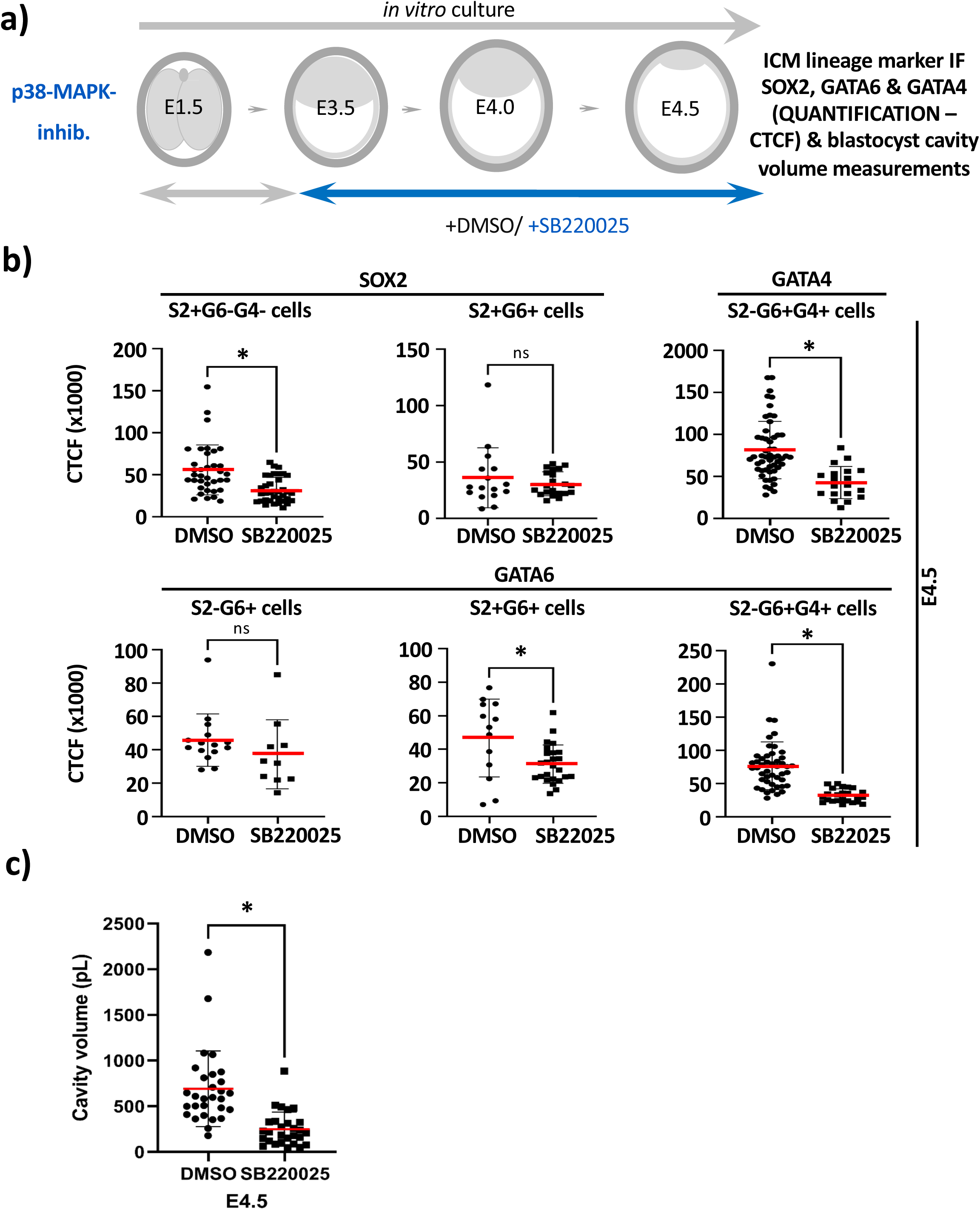
Quantification of average per blastocyst ICM cell expression levels of lineage protein marker expression, and blastocyst cavity volume, after p38-MAPKi treatment (E3.5-E4.0/E4.5; SOX2, GATA6 & GATA4): **a)** Experimental scheme of p38-MAPKi (SB220025 – 20μM) treatment and assay points during blastocyst maturation period (E3.5-E4.5). **b)** Average quantified ICM cell lineage marker expression (SOX2, GATA6 & GATA4; CTCF) in blastocysts cultured from E3.5-E4.5 under control (DMSO) and p38-MAPKi-treated conditions, according to indicated co-expression status; arithmetic means (red bars), standard deviations and statistically significant differences are highlighted (for normal or non-normally distributed data, unpaired T-test or Mann-Whitney tests were employed, respectively; *p<0.05, see supplementary data/stats Excel tables). **c)** Measured blastocyst cavity volumes in i blastocysts cultured from E3.5-E4.5 under control (DMSO) and p38-MAPKi-treated conditions (pL); arithmetic means (red bars), standard deviations and statistically significant differences are highlighted (for the non-normally distributed data, a Mann-Whitney test was employed. *p<0.05; see supplementary data/stats Excel tables).

**Fig. S7:**
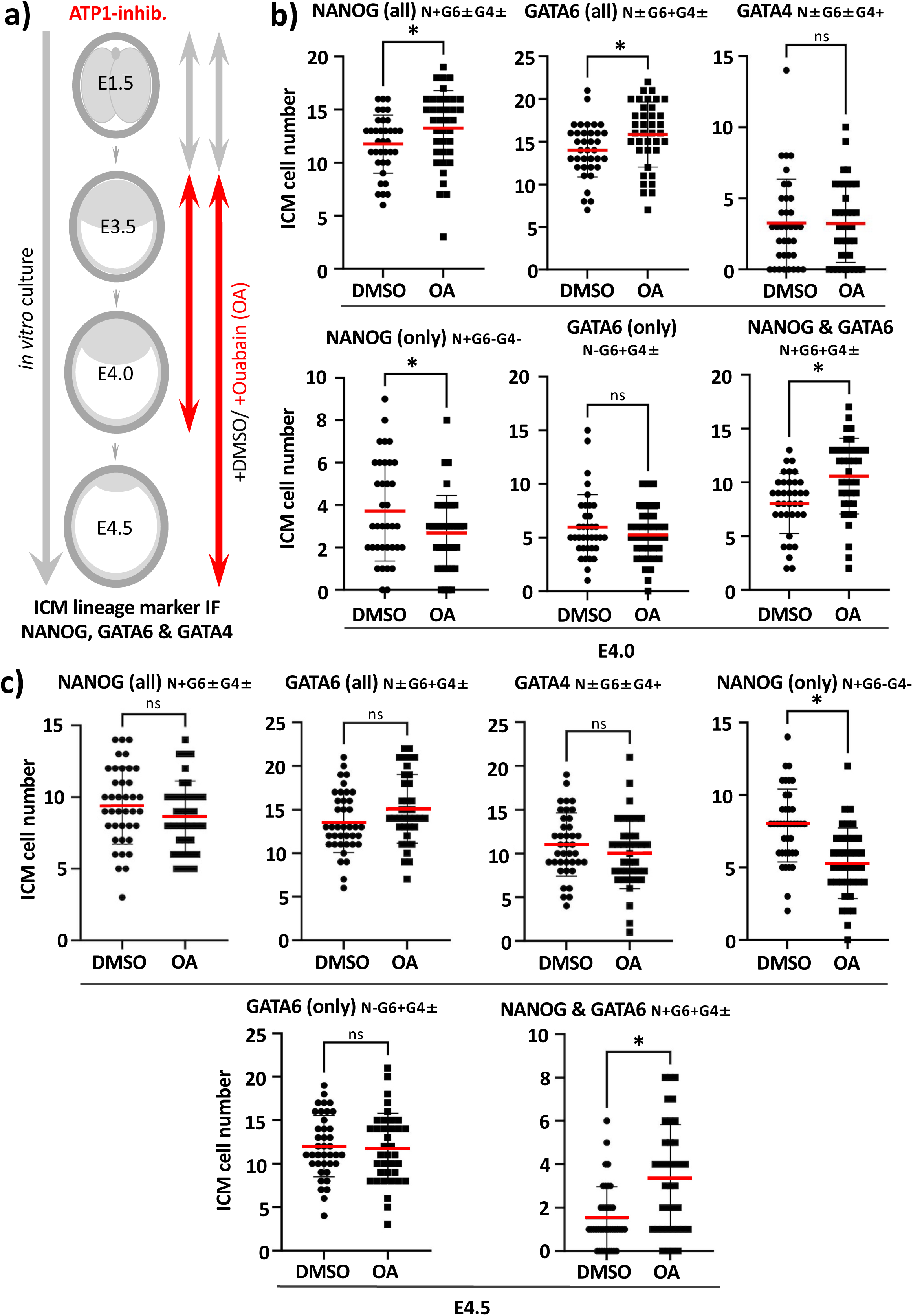
Ouabain treatment during blastocyst maturation (E3.5-E4.0/E4.5) causes impaired numbers of specified EPI from uncommitted ICM progenitors without affecting PrE differentiation (IF: NANOG, GATA6 & GATA4 – average cell number population data): **a)** Experimental scheme of Ouabain (OA) treatment and assay points during blastocyst maturation period (E3.5-E4.5). **b)** Average numbers of total NANOG, total GATA6, total GATA4, NANOG only (without GATA6 or GATA4), GATA6 only (without NANOG, ±GATA4) and NANOG & GATA6 co-expressing (±GATA4) ICM cells, per embryo, assayed at the mid-blastocyst (E4.0) stage under control (DMSO) or OA treatment conditions. **c)** as in b) but assayed at the late-blastocyst (E4.5) stage under control (DMSO) or OA treatment conditions. b) & c); arithmetic means (red bars), standard deviations and statistically significant differences are highlighted (for normal or non-normally distributed data, unpaired T-test or Mann-Whitney tests were employed, respectively; *p<0.05, see supplementary data/stats Excel tables).

**Fig. S8:**
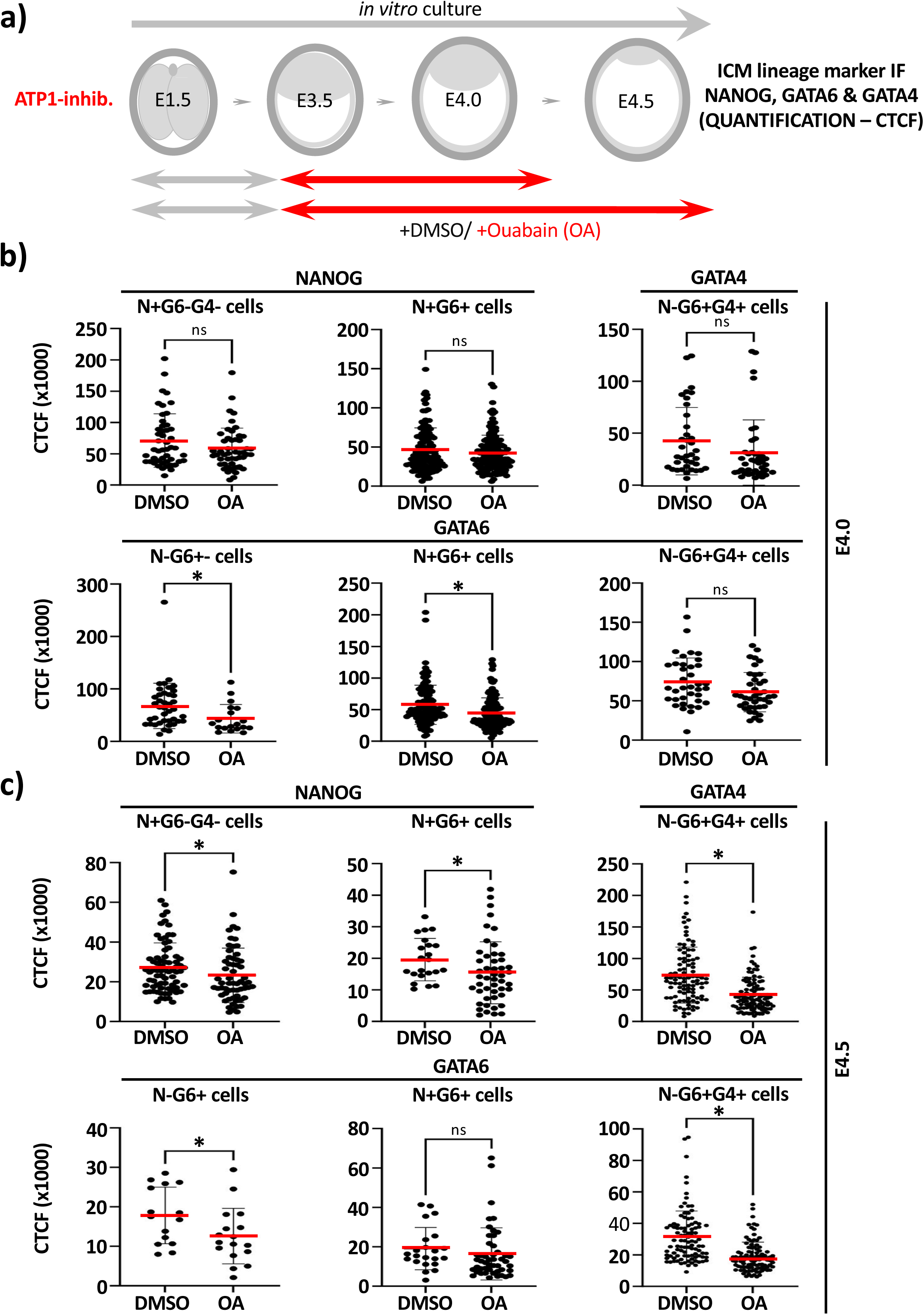
Quantification of average per blastocyst ICM cell expression levels of lineage protein marker expression after Ouabain treatment (E3.5-E4.0/E4.5; NANOG, GATA6 & GATA4): **a)** Experimental scheme of Ouabain (OA) treatment and assay points during blastocyst maturation period (E3.5-E4.5). **b)** Average quantified ICM cell lineage marker expression (NANOG, GATA6 & GATA4; CTCF) in blastocysts cultured from E3.5-E4.0 under control (DMSO) and OA-treated conditions, according to indicated co-expression status. **c)** as in b) for blastocysts cultured from E3.5-E4.5. b) & c); arithmetic means (red bars), standard deviations and statistically significant differences are highlighted (for normal or non-normally distributed data, unpaired T-test or Mann-Whitney tests were employed, respectively; *p<0.05, see supplementary data/stats Excel tables).

**Fig. S9:**
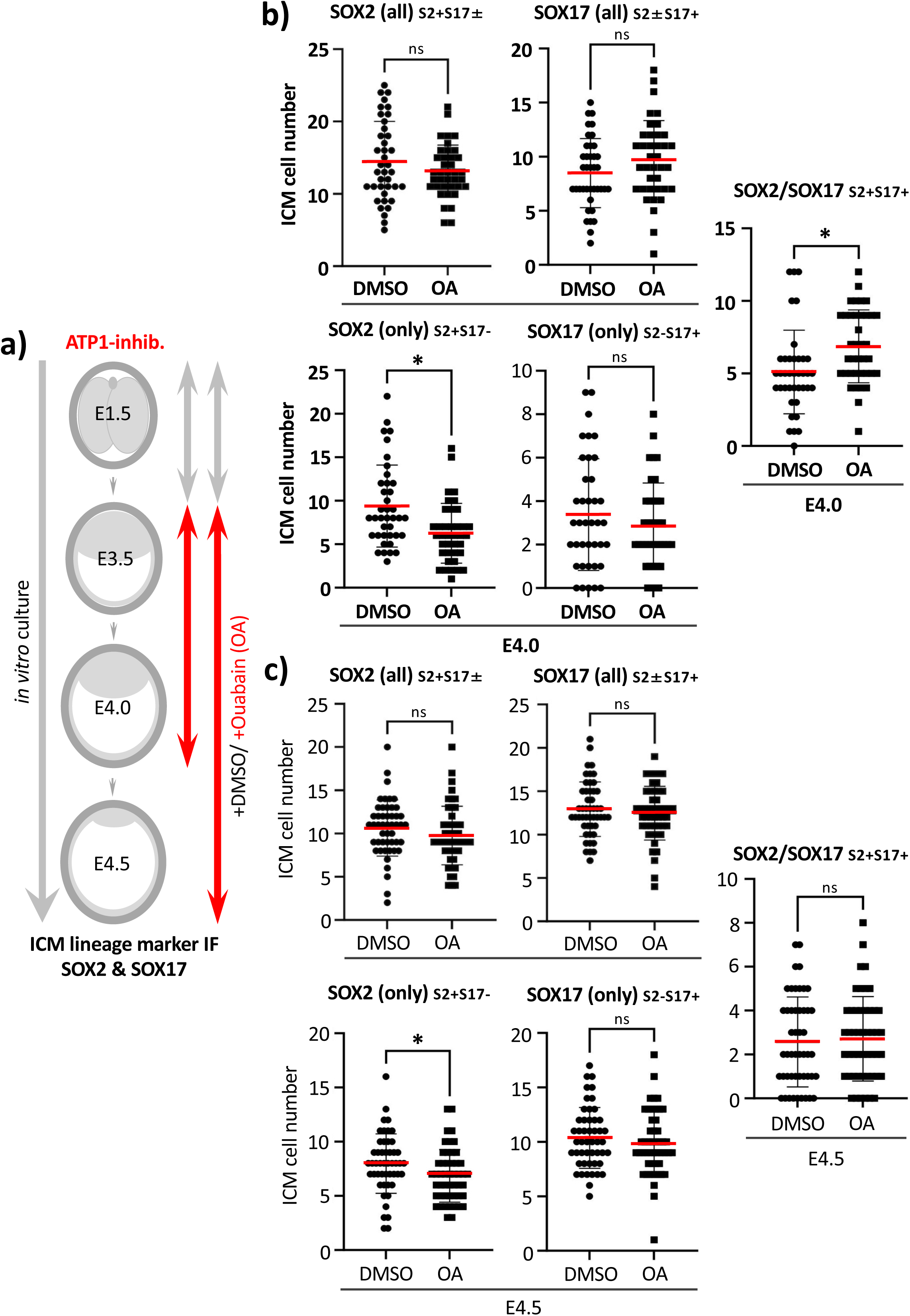
Ouabain treatment during blastocyst maturation (E3.5-E4.0/E4.5) impairs numbers of specified EPI without affecting PrE differentiation (IF: SOX2 & SOX17 – average cell number population data): **a)** Experimental scheme of Ouabain (OA) treatment and assay points during blastocyst maturation period (E3.5-E4.5). **b)** Average numbers of total SOX2, total SOX17, SOX2 only (without SOX17), SOX17 only (without SOX2) and SOX2 & SOX17 co-expressing ICM cells, per embryo, assayed at the mid-blastocyst (E4.0) stage under control (DMSO) or OA treatment conditions. **c)** as in b) but assayed at the late-blastocyst (E4.5) stage under control (DMSO) or OA treatment conditions. b) & c); arithmetic means (red bars), standard deviations and statistically significant differences are highlighted (for normal or non-normally distributed data, unpaired T-test or Mann-Whitney tests were employed, respectively; *p<0.05, see supplementary data/stats Excel tables).

**Fig. S10:**
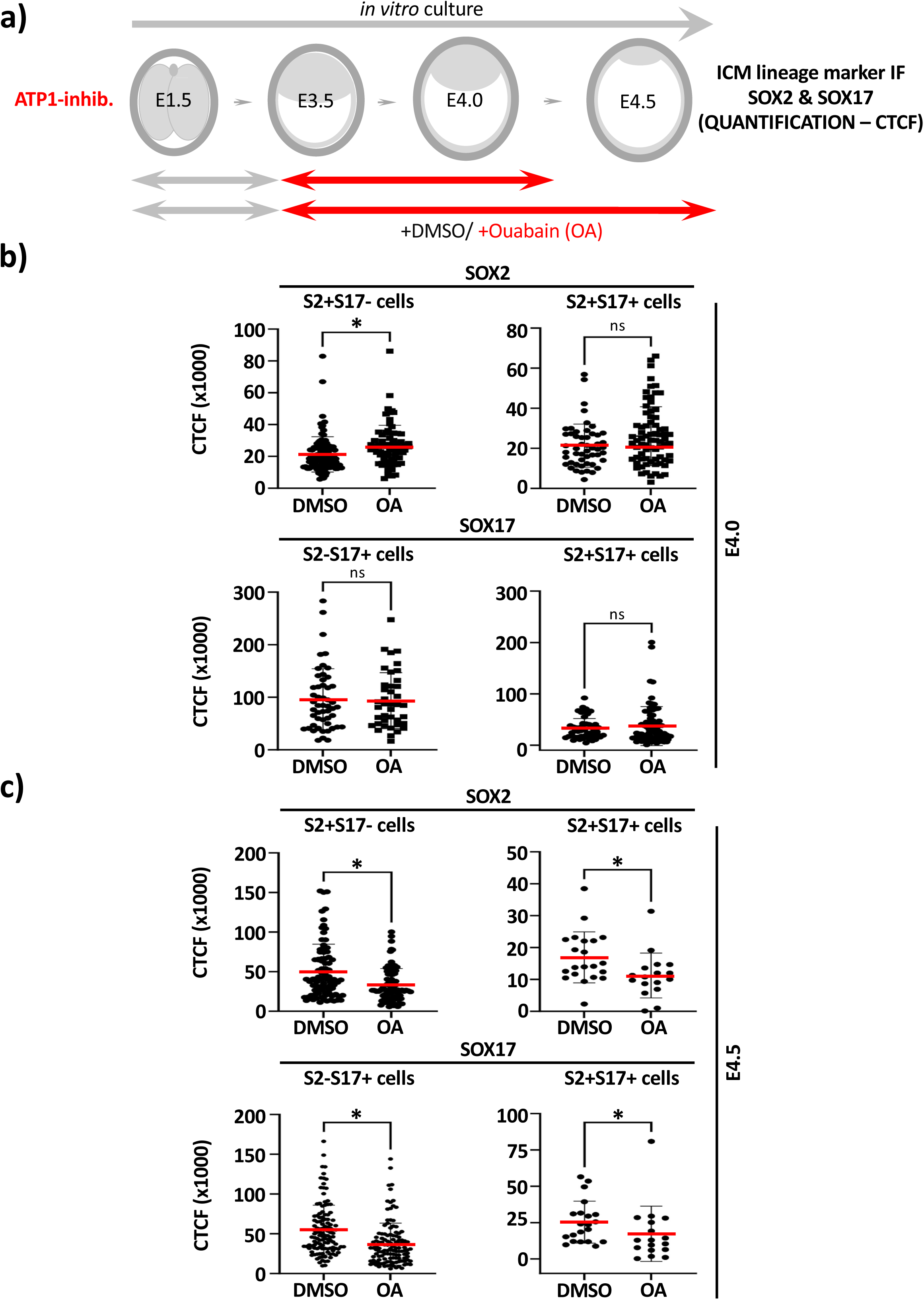
Quantification of average per blastocyst ICM cell expression levels of lineage protein marker expression after Ouabain treatment (E3.5-E4.0/E4.5; SOX2 & SOX17): **a)** Experimental scheme of Ouabain (OA) treatment and assay points during blastocyst maturation period (E3.5-E4.5). **b)** Average quantified ICM cell lineage marker expression (SOX2 & SOX17; CTCF) in blastocysts cultured from E3.5-E4.0 under control (DMSO) and OA-treated conditions, according to co-expression status. **c)** as in b) for blastocysts cultured from E3.5-E4.5. b) & c); arithmetic means (red bars), standard deviations and statistically significant differences are highlighted (for normal or non-normally distributed data, unpaired T-test or Mann-Whitney tests were employed, respectively; *p<0.05, see supplementary data/stats Excel tables).

**Fig. S11:**
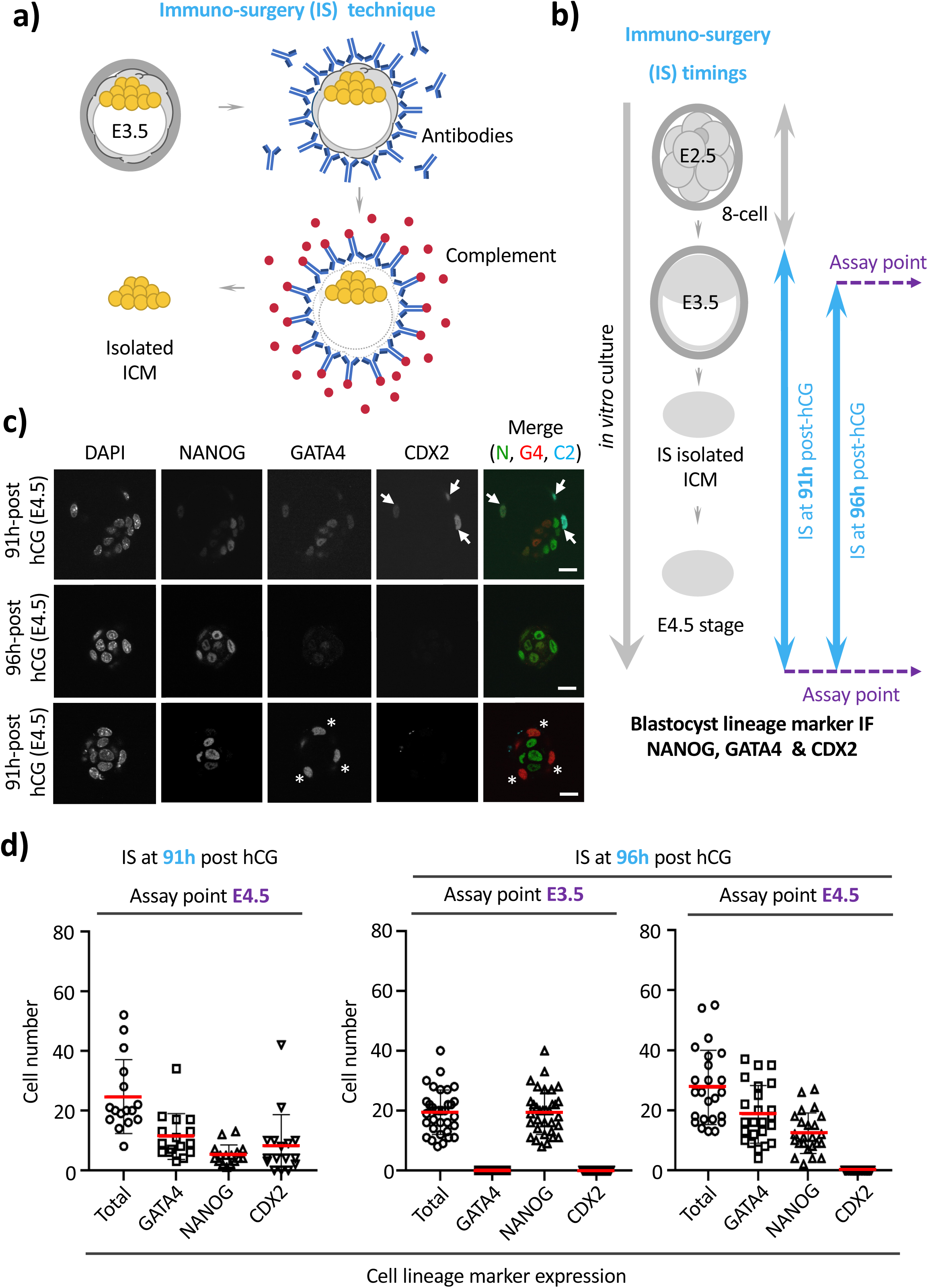
Developmental timing and *in vitro* culture of immuno-surgical isolated (IS) blastocyst ICMs determines whether an outer layer of reconstituted TE or specified PrE is subsequently formed: **a)** Technical IS scheme; *zona*-less early-blastocyst (E3.5) stage embryos were incubated with mouse antibodies, followed by guinea-pig complement to induce complete lysis of outer TE and yield isolated ICMs that can be further *in vitro* cultured and IF assayed for blastocyst cell fate marker protein expression. **b)** Experimental scheme detailing early-blastocyst (E3.5) IS ICM isolation at time-points equivalent to 91 and 96 hours post-hCG superovulation injection and subsequent *in vitro* culture and IF staining against NANOG (EPI), GATA4 (PrE) and CDX2 (TE) cell fate markers. **c)** Example confocal IF stained (NANOG - green, GATA4 - red & CDX2 – cyan) single z-section micrographs of cultured IS isolated ICMs from either time point (scale bar – 20μm). **d)** Average numbers of total, GATA4, NANOG and CDX2 expressing cells per IS isolated and *in vitro* cultured ICM; either assayed immediately after IS isolation at E3.5 (96 hours post-hCG) or following 24 hours of culture at E4.5 (after initial isolation at both 91 & 96 hours post-hCG time points); arithmetic means (red bars) and standard deviations are highlighted; see supplementary data Excel/stats tables. Note in panels c) & d) a lack of detectable CDX2 expressing cells immediately post-IS isolation at E3.5 (indicating successful removal of TE) and reconstitution of CDX2 expressing TE cells (highlighted in micrographs with white arrows) in cultured groups isolated at the earlier 91 hour post-hCG time-point versus specification and formation of outer differentiating PrE expressing GATA4 cells (surrounding NANOG positive inner EPI cells; highlighted with white asterisks in micrographs) in those isolated after 96 hours post-hCG (indicating ICM cell fate resolution equivalent to that of intact cultured blastocysts in the latter group).

**Fig. S12:**
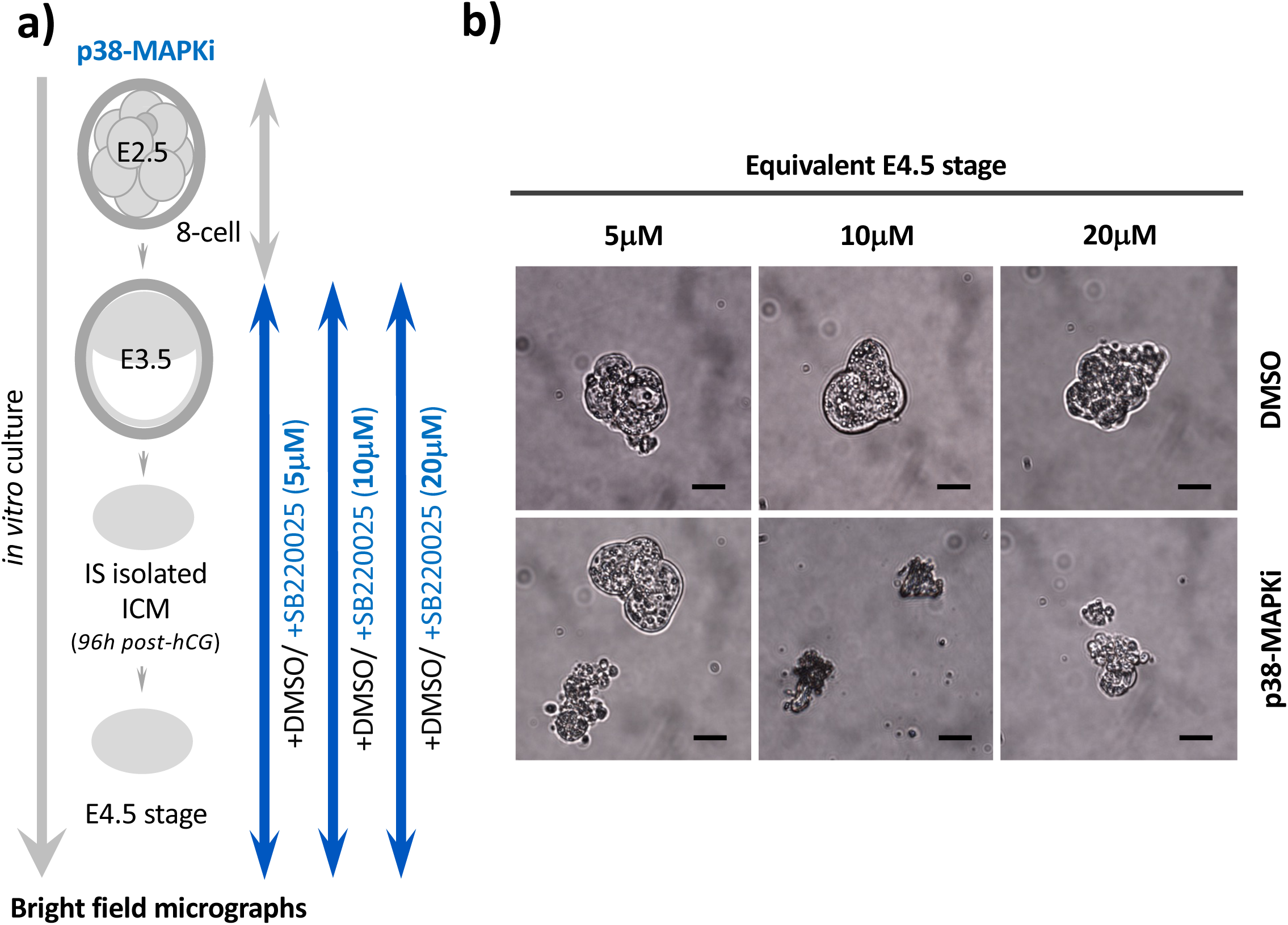
Culture (*in vitro*) of IS isolated ICMs in varying concentrations of p38-MAPKi (SB220025): **a)** Experimental schema by which IS isolated ICMs derived after 96 hours post-hCG (relative to initial superovulation) were cultured for 24 hours in either 5, 10 or 20μM p38-MAPKi conditions (SB220025) or equivalent volumes of DMSO vehicle control and subject to brightfield microscopy to assess ICM viability. **b)** Bright-field micrographs of exemplar control DMSO or p38-MAPKi treated IS isolated ICMs (scale bar - 30μm); note p38-MAPKi induced ICM cytotoxicity at 10 and 20μM, but not 5μM concentrations of SB220025.

**Fig. S13:**
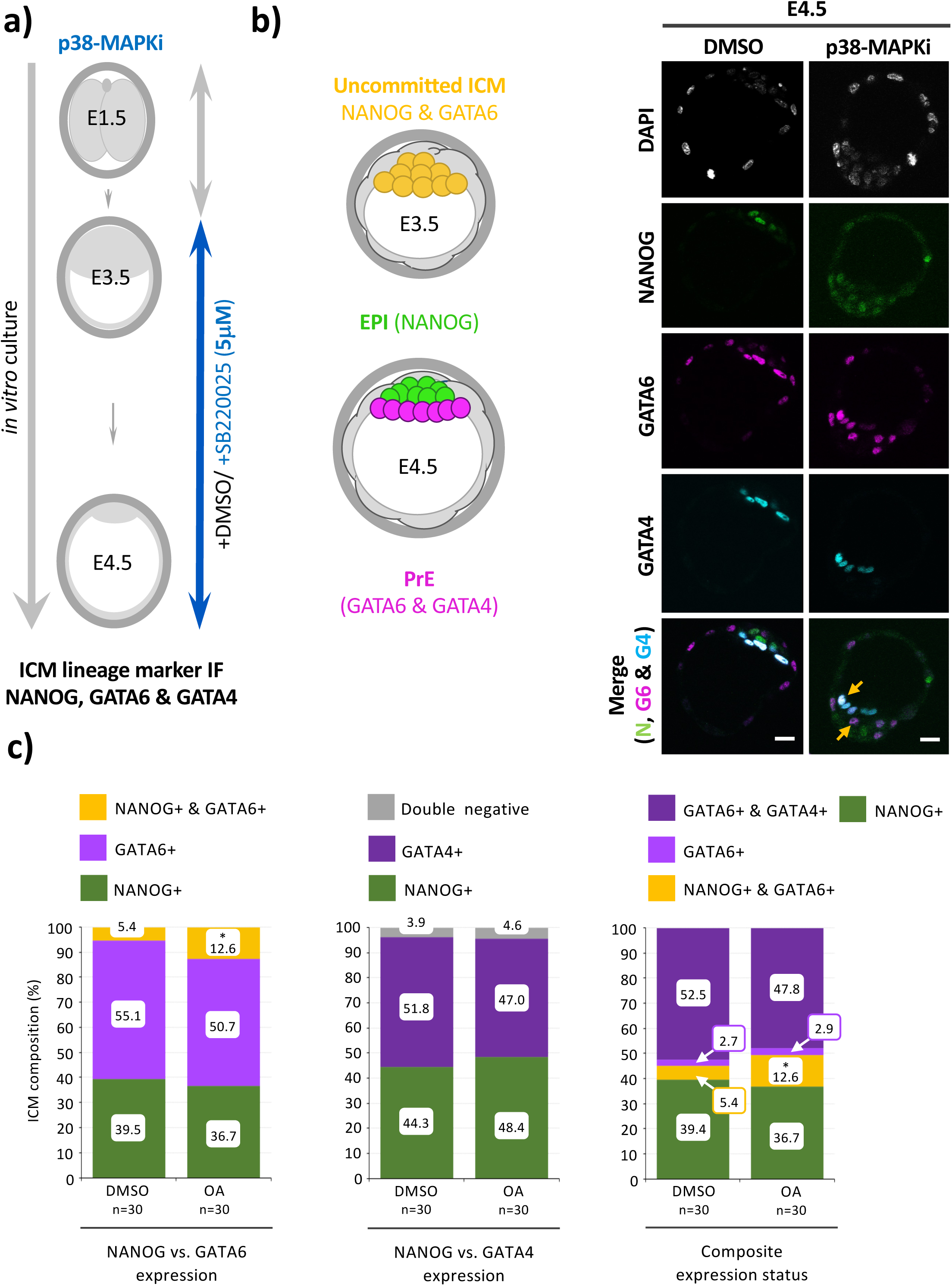
Effect of 5μM SB220025 mediated p38-MAPKi on ICM cell fate derivation during *in vitro* culture of intact mouse blastocysts (E3.5-E4.5 - IF: NANOG, GATA6 & GATA4 – exemplar confocal micrographs and average percentage population data): **a)** Experimental schema by which early-blastocyst (E3.5) stage embryos were cultured for 24 hours to the late-blastocyst (E4.5) stage in either 5μM p38-MAPKi conditions (SB220025) or an equivalent volume of DMSO vehicle control and subject to IF confocal microscopy to assay ICM cell fate marker protein (NANOG, GATA6 & GATA4). **b)** Left - Schematic detailing typical expression of ICM lineage marker proteins in uncommitted early-blastocyst (E3.5) stage (NANOG & GATA6 co-expression - yellow) and at the late-blastocyst (E4.5) stage (EPI – NANOG alone – green, PrE – GATA6 & GATA4 in the absence of NANOG - magenta). Right - Exemplar confocal single z-section micrographs of control (DMSO) or p38-MAPKi treated blastocysts IF assayed for combined NANOG, GATA6 & GATA4 expression at E4.5 (scale bar - 20μm, arrows denote uncommitted ICM cells co-expressing NANOG & GATA6). **c)** Average cellular percentage ICM composition of control (DMSO) or p38-MAPKi treated blastocysts IF assayed for cell lineage marker expression (panel b); delineated as i. uncommitted (co-expressing NANOG & GATA6), PrE (expressing GATA6 without NANOG) and EPI (expressing NANOG without GATA6) cells (left panel), ii. PrE (expressing GATA4 without NANOG) and EPI (expressing NANOG without GATA4) cells (central panel), or iii. uncommitted (co-expressing NANOG & GATA6), PrE (expressing GATA6 alone or GATA6 plus GATA4 without NANOG) and EPI (expressing NANOG without GATA6 or GATA4) cells (right panel). Statistically significant differences highlighted (Z-test *p<0.05, n-numbers reflect total blastocysts assayed; see supplementary data/stats Excel tables). *See Fig. S14 for average cell number population data/per assayed embryo.* Note, p38-MAPKi results in increased incidence of uncommitted PrE progenitor cells (revealed by co-expression of NANOG and GATA6 – see panel b) highlighted by yellow arrows).

**Fig. S14:**
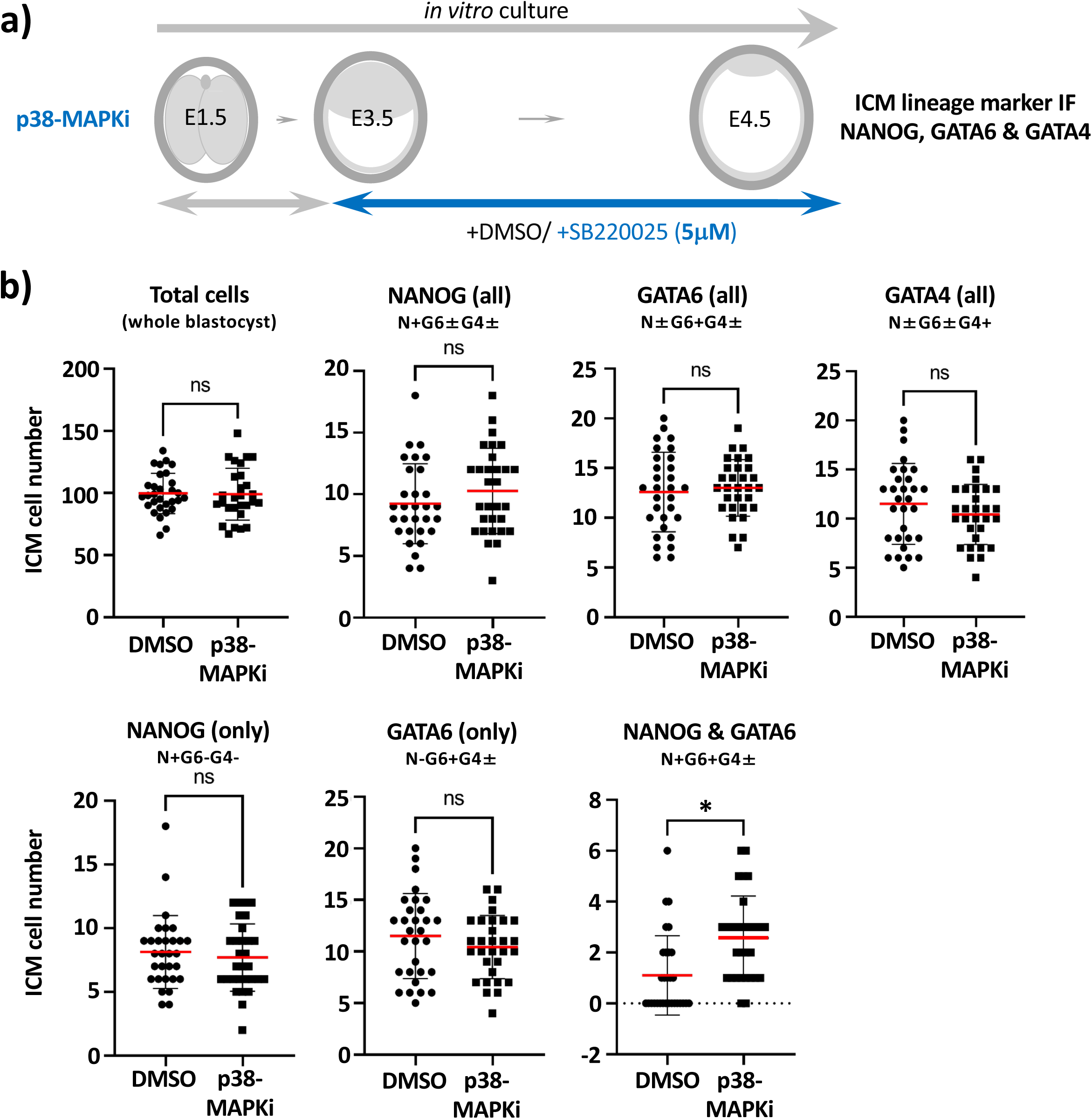
Effect of 5μM SB220025 mediated p38-MAPKi on ICM cell fate derivation during *in vitro* culture of intact mouse blastocysts (E3.5-E4.5 - IF: NANOG, GATA6 & GATA4 – average cell number population data): **a)** Experimental schema by which early-blastocyst (E3.5) stage embryos were cultured for 24 hours to the late-blastocyst (E4.5) stage in either 5μM p38-MAPKi conditions (SB220025) or equivalent volumes of DMSO vehicle control and subject to IF confocal microscopy to assay ICM cell fate marker protein (NANOG, GATA6 & GATA4). **b)** Average numbers of total blastocyst, total NANOG, total GATA6, total GATA4, NANOG only (without GATA6 or GATA4), GATA6 only (without NANOG, ±GATA4) and NANOG & GATA6 co-expressing (±GATA4) ICM cells, per embryo, assayed at the late-blastocyst (E4.5) stage under control (DMSO) or p38-MAPKi (5μM) treatment conditions; arithmetic means (red bars), standard deviations and statistically significant differences are highlighted (for normal or non-normally distributed data, unpaired T-test or Mann-Whitney tests were employed, respectively; *p<0.05); see supplementary data/stats Excel tables.

**Fig. S15:**
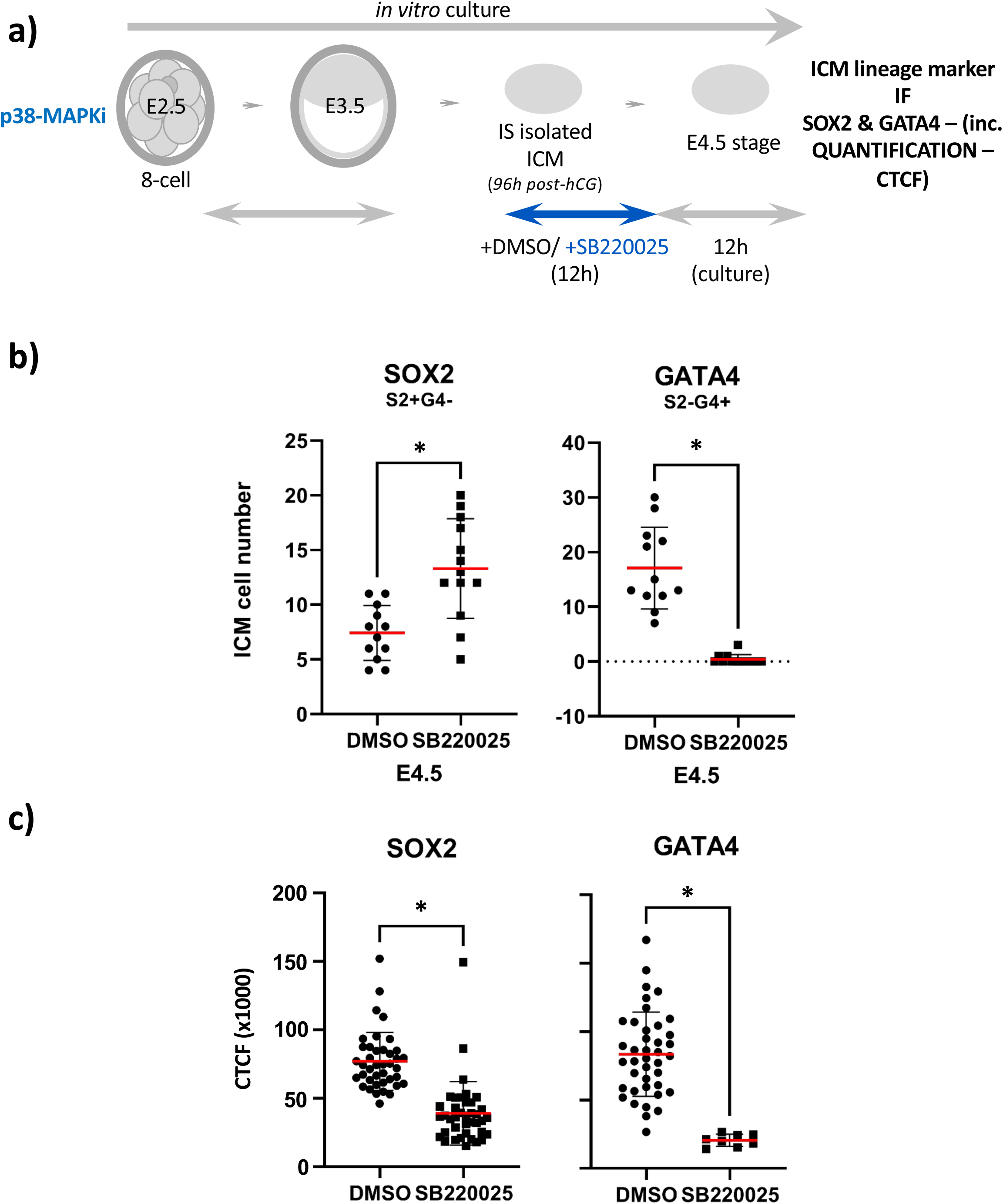
p38-MAPKi treatment of *in vitro* cultured early-blastocyst (E3.5) stage IS isolated ICMs effectively impairs specification and differentiation of the PrE lineage (IF: SOX2, GATA6 & GATA4 – average cell number population data): **a)** Experimental scheme by which early-blastocyst (E3.5 – 96 hours post-hCG superovulation microinjection) stage IS isolated ICMs were cultured for 12 hours under control (DMSO) or p38-MAPKi (5μM SB220025) conditions before transfer into regular growth media for a further 12 hours, until the equivalent late-blastocyst (E4.5) stage, and IF stained for ICM lineage markers. **b)** Average numbers of total SOX2 and total GATA4 expressing (in the absence of the other assayed marker), per IS isolated ICM, assayed at the equivalent late-blastocyst (E4.5) stage after prior control (DMSO) or p38-MAPKi treatment; arithmetic means (red bars), standard deviations and statistically significant differences are highlighted (for normal or non-normally distributed data, unpaired T-test or Mann-Whitney tests were employed, respectively; *p<0.05); see supplementary data/stats Excel tables. **c)** Average quantified ICM cell lineage marker expression (SOX2 & GATA4; CTCF) in IS-isolated ICMs cultured from E3.5-E4.0 under control (DMSO) and p38-MAPKi-treated conditions, and then further cultured until the E4.5 equivalent stage in regular culture media, according to co-expression status; arithmetic means (red bars), standard deviations and statistically significant differences are highlighted (for normal or non-normally distributed data T-test or Mann-Whitney tests were employed, respectively; *p<0.05, see supplementary data/stats Excel tables).

**Fig. S16:**
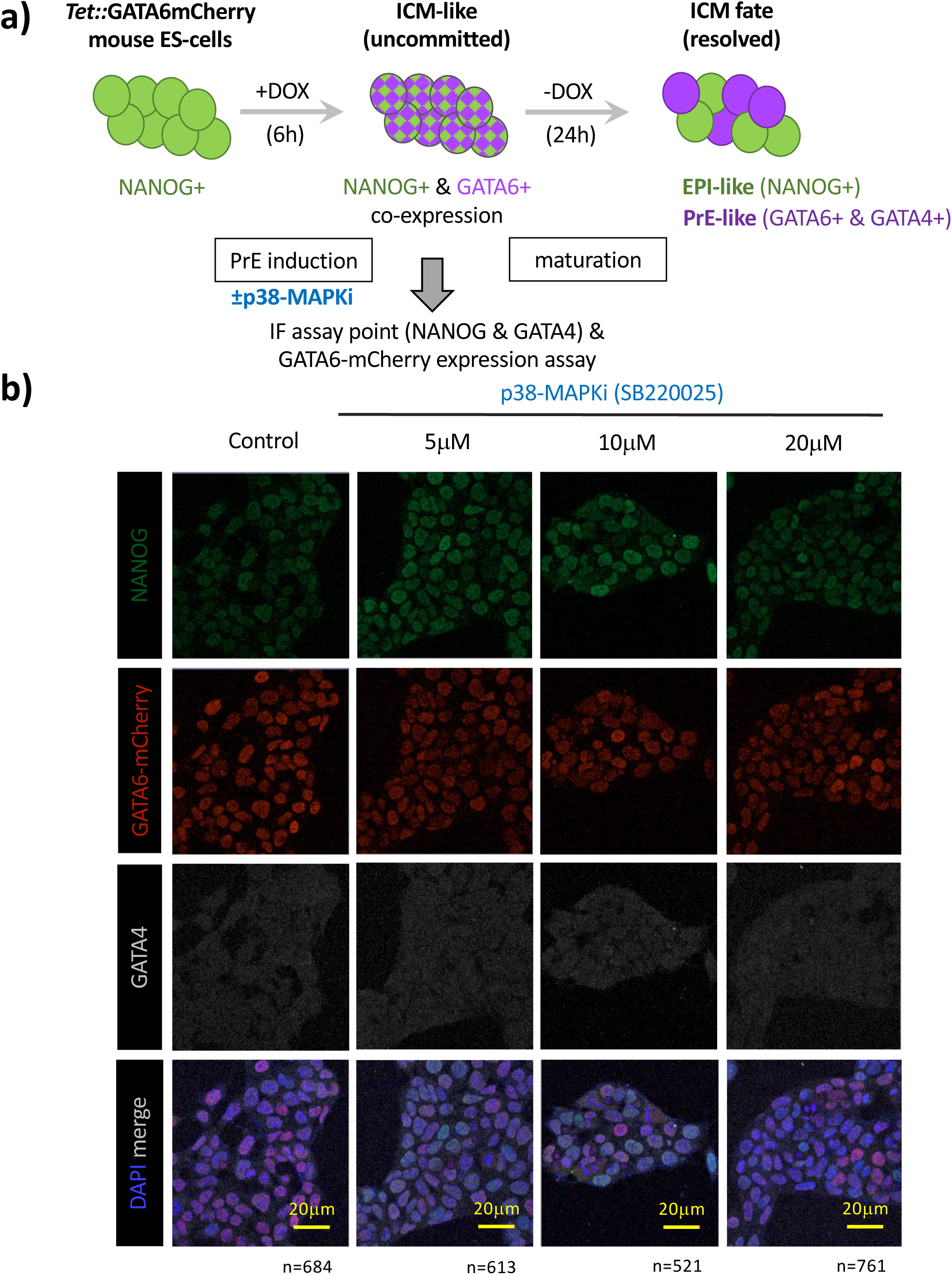
Induction of recombinant GATA6-mCherry expression in a mouse ES-cell model of blastocyst ICM fate derivation under control and p38-MAPKi conditions (IF: NANOG, GATA6 & GATA4 – example confocal micrographs): **a)** Experimental schema by which the mouse ES-cell line *Tet::Gata6mCherry* can be induced by a 6-hour pulse of doxycycline treatment (+DOX 6h) to co-express endogenous NANOG and a recombinant GATA6-mCherry fusion protein (reminiscent of uncommitted early-blastocyst/E3.5 stage ICM (Chazaud et al. 2006)). After a further 24 hours of culture in regular culture media (-DOX 24h), individual cells can mature to express NANOG and endogenous GATA6/GATA4 in a mutually exclusive pattern (Schroter et al. 2015) (akin to ICM lineage segregation in the late blastocyst/E4.5 stage (Chazaud et al. 2006)). Here, ES-cell cultures were exposed during +DOX 6h induction to p38-MAPKi (using SB220025 at 5, 10 or 20μM) or vehicle control (DMSO) and immediately subject to IF expression analysis for the pluripotency marker NANOG or the PrE marker GATA4 (+induced GATA6-mCherry); to confirm the formation of the uncommitted early-blastocyst/E3.5 stage ICM-like stage in control and assaying the effect of p38-MAPKi treatment. **b)** Example confocal micrographs of cell regions of interest cultured under the indicated +DOX 6h conditions, IF stained for endogenous NANOG (green) and GATA4 (grey-scale), plus induced recombinant GATA6-mCherry (red) expression; a full merge image with DAPI counterstain is provided (lower panels). Scale bars - 20μm.

**Fig. S17:**
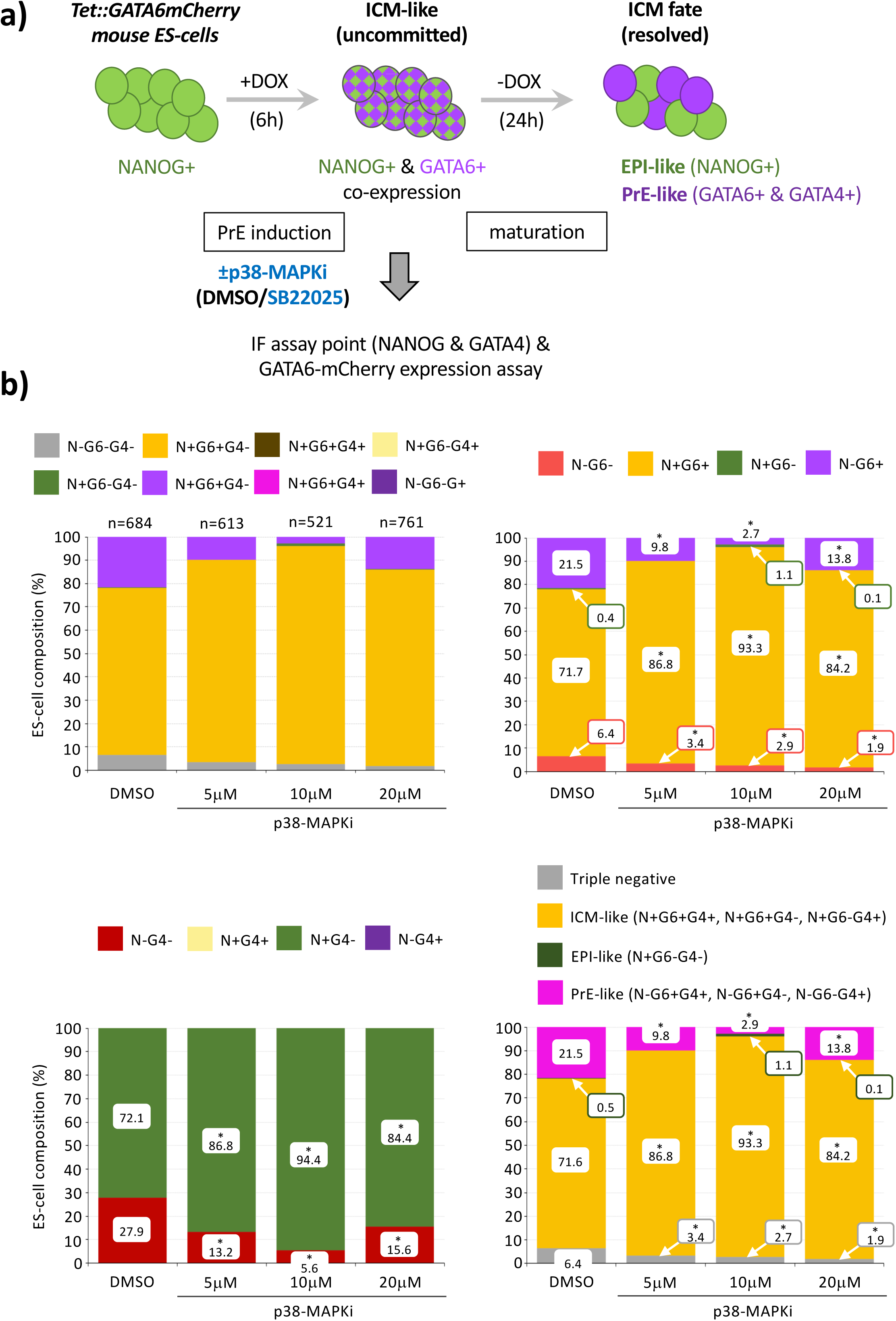
Induction of recombinant GATA6-mCherry expression in a mouse ES-cell model of blastocyst ICM fate derivation under control and p38-MAPKi conditions (IF: NANOG, GATA6 & GATA4 – average percentage population data): **a)** Experimental schema by which the mouse ES-cell line *Tet::Gata6mCherry* can be induced by a 6-hour pulse of doxycycline treatment (+DOX 6h) to co-express endogenous NANOG and a recombinant GATA6-mCherry fusion protein (reminiscent of uncommitted early-blastocyst/E3.5 stage ICM (Chazaud et al. 2006)). After a further 24 hours of culture in regular culture media (-DOX 24h), individual cells can mature to express NANOG and endogenous GATA6/GATA4 in a mutually exclusive pattern (Schroter et al. 2015) (akin to ICM lineage segregation in the late blastocyst/E4.5 stage (Chazaud et al. 2006)). Here, ES-cell cultures were exposed during +DOX 6h induction to p38-MAPKi (using SB220025 at 5, 10 or 20μM) or vehicle control (DMSO) and immediately subject to IF expression analysis for the pluripotency marker NANOG or the PrE marker GATA4 (plus induced GATA6-mCherry); to confirm the formation of the uncommitted early-blastocyst/E3.5 stage ICM like stage in control and assaying the effect of p38-MAPKi treatment. **b)** Average cellular percentage composition of control (DMSO) or p38-MAPKi treated induced (+DOX 6h) *Tet::Gata6mCherry* ES-cells and IF assayed (*as shown in Fig. S16*) for denoted marker protein expression; i. all possible cell lineage marker expression combinations (upper-left panel; +/-N = NANOG, +/-G6 = GATA6 & +/-G4 = GATA4), ii. compared (co-)expression status of NANOG & GATA6 (upper-right panel), iii. compared (co-)expression status of NANOG & GATA4 (lower-left panel) and iv. classified as blastocyst PrE-like, EPI-like or ICM-like (uncommitted (Chazaud et al. 2006)) cell lineages or triple-negative cells (lacking detectable marker protein expression; lower-right panel). Individual percentage ICM contributions and statistically significant differences against the control condition are highlighted (Z-test, *p<0.05; and n-numbers denote total cells assayed).

**Fig. S18:**
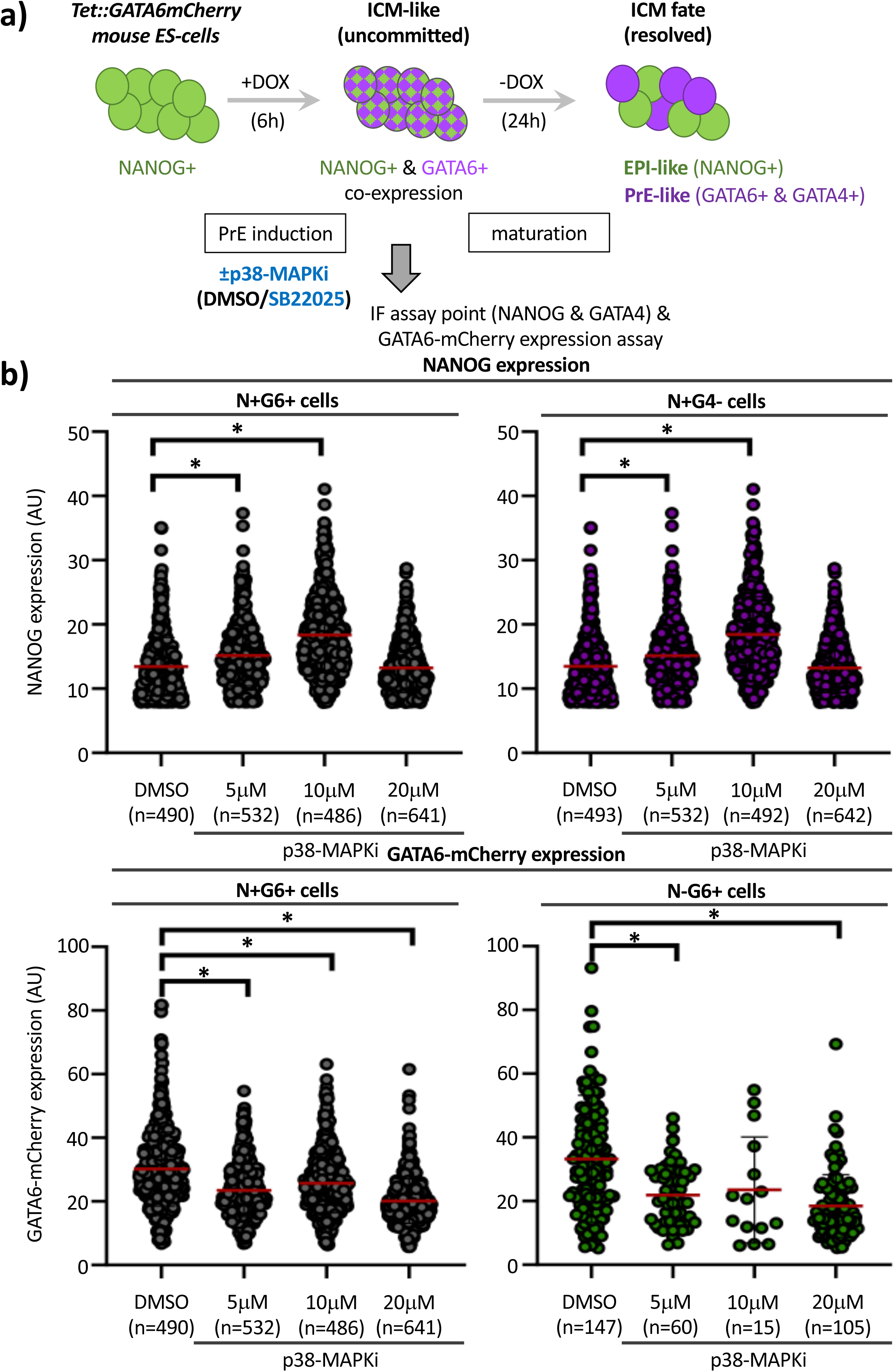
Induction of recombinant GATA6-mCherry expression in a mouse ES-cell model of blastocyst ICM fate derivation under control and p38-MAPKi conditions (IF: NANOG, GATA6 & GATA4 – average quantified, per cell, lineage marker protein expression): **a)** Experimental schema by which the mouse ES-cell line *Tet::Gata6mCherry* can be induced by a 6-hour pulse of doxycycline treatment (+DOX 6h) to co-express endogenous NANOG and recombinant GATA6-mCherry fusion protein (reminiscent of uncommitted early-blastocyst/E3.5 stage ICM (Chazaud et al. 2006)). After a further 24 hours of culture in regular culture media (-DOX 24h), individual cells can mature to express NANOG and endogenous GATA6/GATA4 in a mutually exclusive pattern (Schroter et al. 2015) (akin to ICM lineage segregation in the late blastocyst/E4.5 stage (Chazaud et al. 2006)). Here, ES-cell cultures were exposed during +DOX 6h induction to p38-MAPKi (using SB220025 at 5, 10 or 20μM) or vehicle control (DMSO) and immediately subject to IF expression analysis for the pluripotency marker NANOG or the PrE marker GATA4 (+induced GATA6-mCherry); to confirm the formation of the uncommitted early-blastocyst/E3.5 stage ICM like stage in control and assaying the effect of p38-MAPKi treatment. **b)** Average quantified cell lineage marker expression (endogenous NANOG & recombinant GATA6-mCherry; AU), per assayed +DOX 6h induced ES-cell under control (DMSO) and p38-MAPKi inhibited (SB22025 - 5, 10 & 20μM) conditions; delineated by combined lineage marker expression status (+/-N = NANOG, +/-G6 = GATA6 & +/-G4 = GATA4); arithmetic means (red bars), standard deviations and statistically significant differences between control (DMSO) and individual p38-MAPKi treatments are highlighted (Z-test, *p<0.05; n-numbers reflect total cells from each group assayed).

**Fig. S19:**
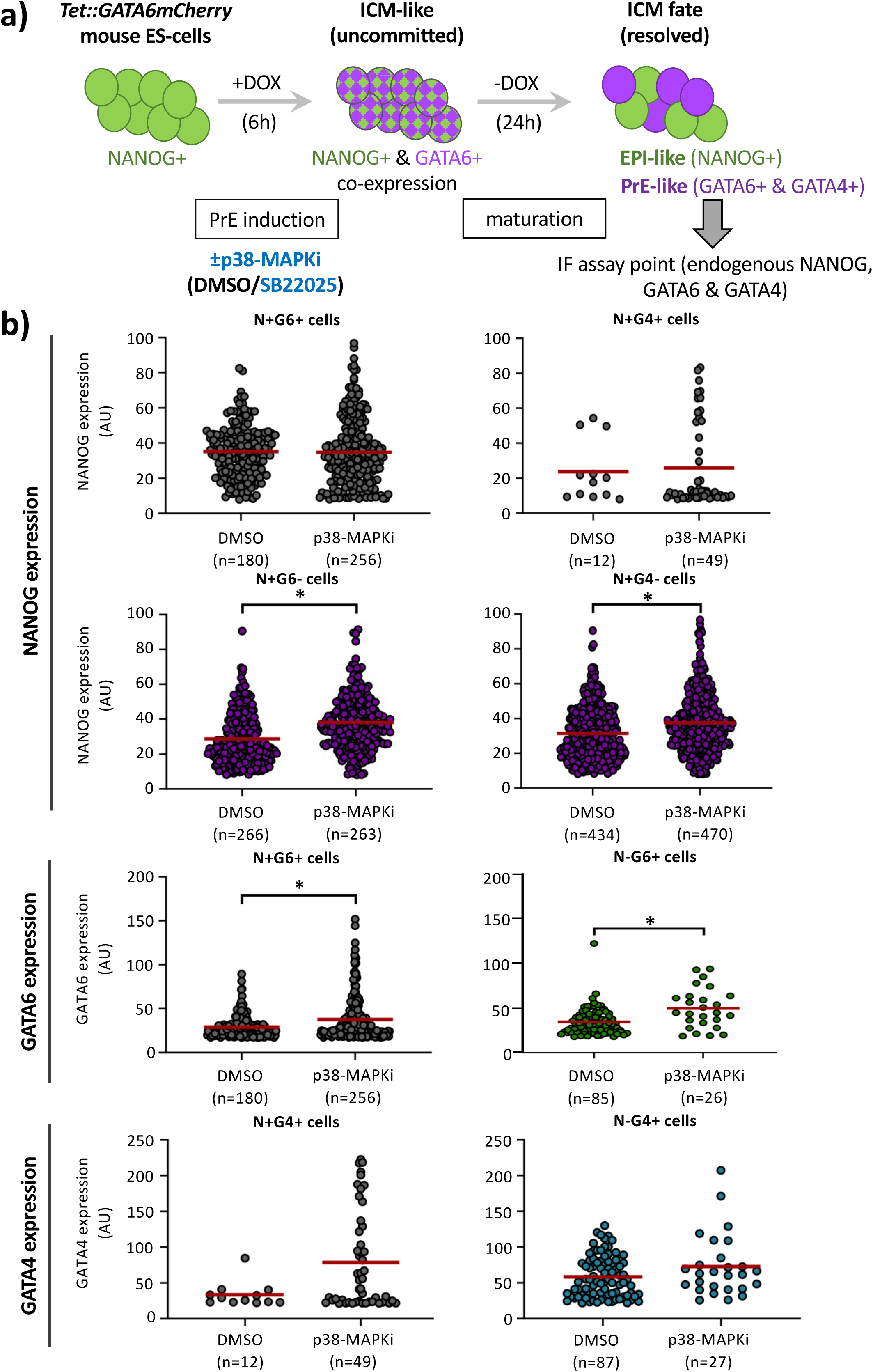
p38-MAPKi impairs PrE-like specification and differentiation in a 2D recombinant GATA6-inducible mouse ES-cell model (IF: NANOG, GATA6 & GATA4 – per cell average quantified lineage marker protein expression): **a)** The mouse ES-cell line *Tet::Gata6mCherry* is be induced by a 6-hour pulse of doxycycline treatment (+DOX 6h) to co-express endogenous NANOG and recombinant GATA6-mCherry fusion protein (reminiscent of uncommitted early-blastocyst/E3.5 stage ICM (Chazaud et al. 2006)). After a further 24 hours of culture in regular culture media (-DOX 24h), individual cells mature to express NANOG and endogenous GATA6/GATA4 in a mutually exclusive pattern (Schroter et al. 2015) (akin to ICM lineage segregation in the late blastocyst/E4.5 stage (Chazaud et al. 2006)). Hence, providing a 2D ES-cell culture model of blastocyst ICM fate derivation (Schroter et al. 2015), against which the effect of p38-MAPKi treatment can be assayed. Accordingly, ES-cell cultures were exposed during +DOX 6h induction to p38-MAPKi (SB220025; 10μM) or vehicle control (DMSO) before transfer into regular culture -DOX 24h (serum + LIF) conditions for 24 hours, followed by fixation and cell lineage marker protein assay (triple IF against NANOG, GATA6 & GATA4). **b)** Average quantified cell lineage marker expression (endogenous NANOG, GATA6 & GATA4; AU) per assayed +DOX 6h induced ES-cell, under control (DMSO) and p38-MAPKi inhibited conditions (and cultured for a further 24 hours), delineated by combined lineage marker expression status (+/-N = NANOG, +/-G6 = GATA6 & +/-G4 = GATA4); arithmetic means (red bars) and statistically significant differences between control (DMSO) and individual p38-MAPKi treatments are highlighted (Z-test, *p<0.05; n-numbers denote total cells assayed per indicated group).

**Fig. S20:**
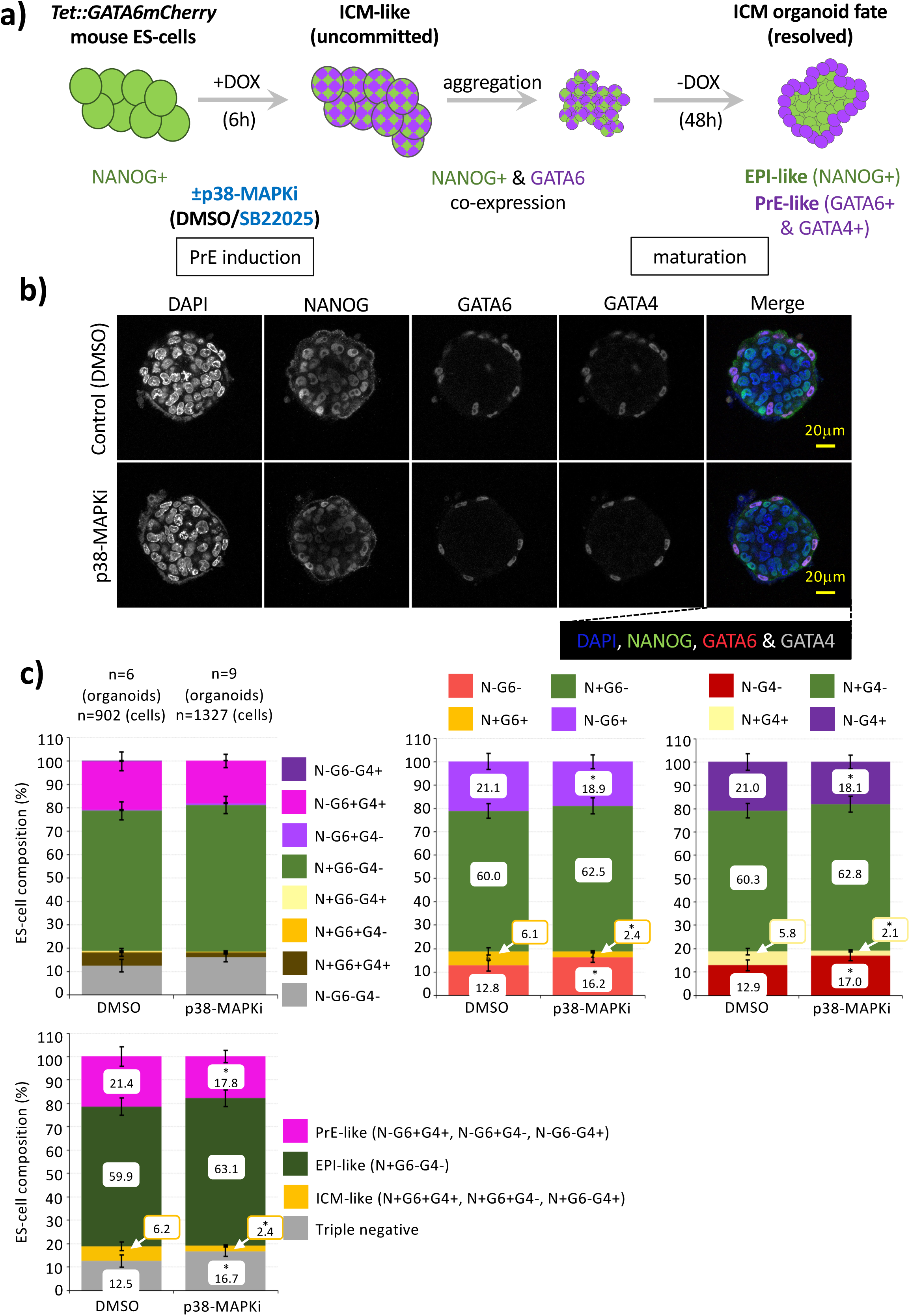
Effect of p38-MAPKi on the derivation of ICM-like cell fate in a mouse ICM organoid model (IF: NANOG, GATA6 & GATA4 – exemplar confocal micrographs & average percentage population data): **a)** Under control conditions, the mouse ES-cell line *Tet::Gata6mCherry* can be induced by a 6-hour pulse of doxycycline treatment (+DOX 6h) to co-express endogenous NANOG and a recombinant GATA6-mCherry fusion protein (Schroter et al. 2015) (reminiscent of uncommitted early-blastocyst/E3.5 stage ICM (Chazaud et al. 2006)). Following induction, 50 individual cells can be collated in concave tissue culture wells and cultured under regular conditions (serum + LIF) for 48 hours (-DOX 48h) to generate aggregate ICM-like organoids comprising a NANOG-expressing (EPI-like) inner core of cells surrounded by an outer-monolayer of endogenous GATA6/4 positive cells (Mathew et al. 2019) (akin to ICM lineage segregation in the late blastocyst/E4.5 stage (Chazaud et al. 2006)); as assayable by IF. Hence, providing a 3D ES-cell-based ICM organoid model of blastocyst ICM fate derivation (as previously described (Mathew et al. 2019)), against which the effect of p38-MAPKi treatment can be tested. The effect on cell lineage derivation of p38-MAPKi (SB220025 10μM) during the +DOX 6h induction period was assayed in ICM organoids generated as described. **b)** Example confocal micrographs of mouse ES-cell ICM organoids derived under the indicated +DOX 6h induction conditions, aggregated and further cultured for 48 hours (-DOX 48h) and IF stained for endogenous NANOG, GATA6 and GATA4 expression (+DAPI, grey-scale); a full pseudo-coloured merge image is provided (right-hand panels). Scale bars - 20μm. **c)** Average cellular percentage ICM organoid composition of control (DMSO) or p38-MAPKi treated induced (+DOX 6h) *Tet::Gata6mCherry* ES-cells used to derive assayed ICM organoids as IF assayed (panel b) for denoted marker protein expression; i. all possible cell lineage marker expression combinations (upper-left panel; +/-N = NANOG, +/-G6 = GATA6 & +/-G4 = GATA4), ii. compared (co-)expression status of NANOG & GATA6 (upper-mid panel), iii. compared (co-)expression status of NANOG & GATA4 (upper-right panel) and iv. classified as blastocyst PrE-like, EPI-like or ICM-like (uncommitted (Chazaud et al. 2006)) cell lineages or triple negative cells (lacking detectable marker protein expression; lower right panel). Individual percentage ICM contributions and statistically significant differences are highlighted (Z-test, *p<0.05; n-numbers denote total ICM organoids and constituent cell numbers assayed).

**Fig. S21:**
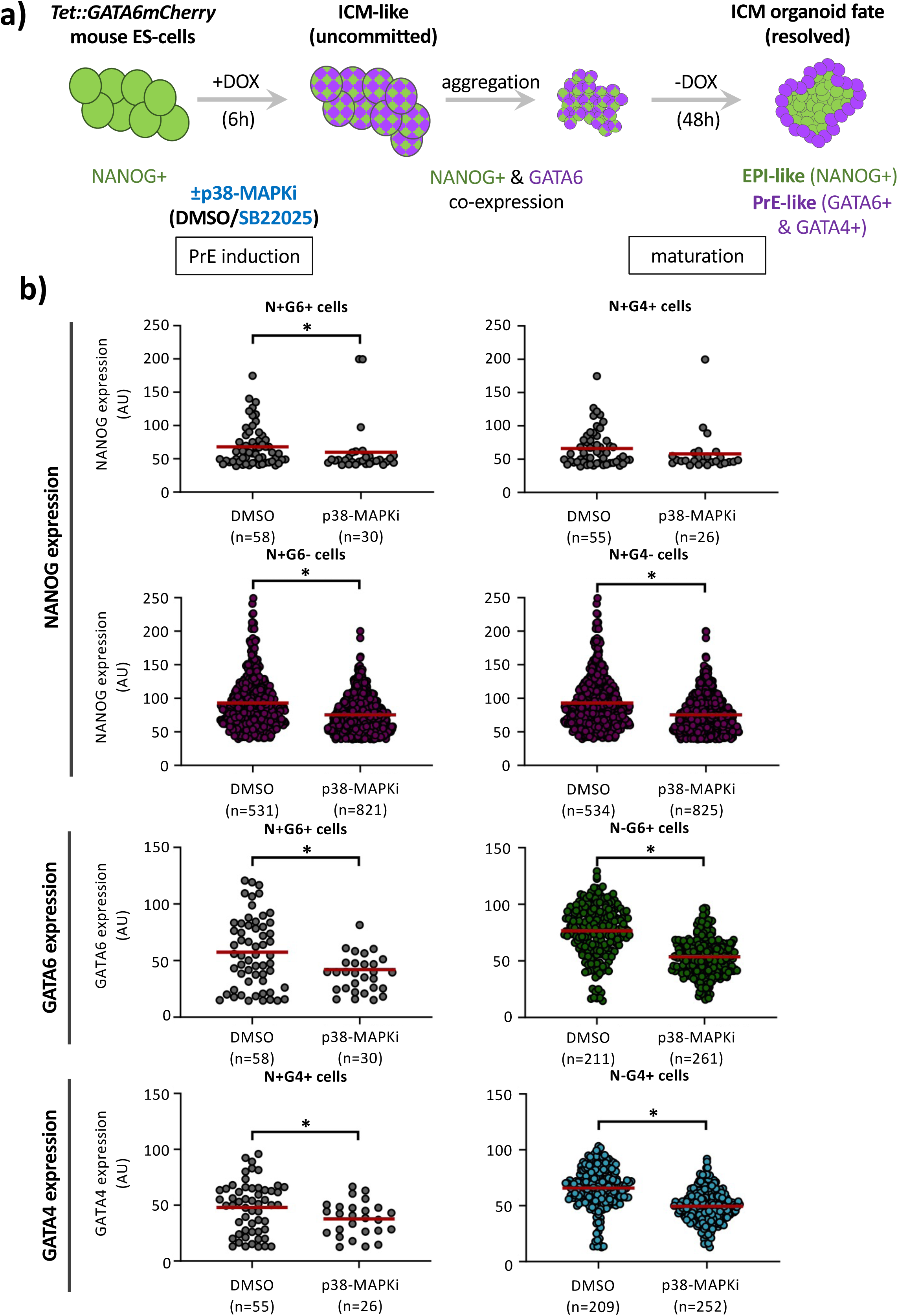
Effect of p38-MAPKi on the derivation of ICM-like cell fate in a 3D recombinant GATA6-inducible mouse ES-cell ICM organoid model (IF: NANOG, GATA6 & GATA4 – per cell average quantified lineage marker protein expression): **a)** Under control conditions, the mouse ES-cell line *Tet::Gata6mCherry* can be induced by a 6-hour pulse of doxycycline treatment (+DOX 6h) to co-express endogenous NANOG and a recombinant GATA6-mCherry fusion protein (Schroter et al. 2015) (reminiscent of uncommitted early-blastocyst/E3.5 stage ICM (Chazaud et al. 2006)). Following induction, 50 individual cells can be collated in concave tissue culture wells and cultured under regular conditions (serum + LIF) for 48 hours (-DOX 48h) to generate aggregate ICM-like organoids comprising a NANOG expressing (EPI-like) inner core of cells surrounding an outer-monolayer of endogenous GATA6/4 positive cells (Mathew et al. 2019) (akin to ICM lineage segregation in the late blastocyst/E4.5 stage (Chazaud et al. 2006)); as assayable by IF. Hence providing a 3D ES-cell based ICM organoid model of blastocyst ICM fate derivation (as previously described (Mathew et al. 2019)), against which the effect of p38-MAPKi treatment can be tested. Accordingly, the effect on cell lineage derivation of p38-MAPKi (SB220025; 10μM) during the +DOX 6h induction period was assayed in ICM organoids generated as described. **b)** Average quantified cell lineage marker expression (endogenous NANOG, GATA6 & GATA4; AU), per assayed ICM organoid cell derived from control (DMSO) or p38-MAPKi treated induced (+DOX 6h) *Tet::Gata6mCherry* ES-cells, delineated by combined lineage marker expression status (+/-N = NANOG, +/-G6 = GATA6 & +/-G4 = GATA4); arithmetic means (red bars), standard deviations and statistically significant differences between control (DMSO) and individual p38-MAPKi treatments are highlighted (Z-test, *p<0.05; n-numbers denote total ICM organoids in each group).

## REFERENCES

Artus J, Piliszek A, Hadjantonakis AK. 2011. The primitive endoderm lineage of the mouse blastocyst: sequential transcription factor activation and regulation of differentiation by Sox17. Dev Biol 350: 393–404.

Avilion AA, Nicolis SK, Pevny LH, Perez L, Vivian N, Lovell-Badge R. 2003. Multipotent cell lineages in early mouse development depend on SOX2 function. Genes Dev 17: 126–140.

Bagnat M, Cheung ID, Mostov KE, Stainier DY. 2007. Genetic control of single lumen formation in the zebrafish gut. Nat Cell Biol 9: 954–960.

Bora P, Gahurova L, Hauserova A, Stiborova M, Collier R, Potesil D, Zdrahal Z, Bruce AW. 2021a. DDX21 is a p38-MAPK-sensitive nucleolar protein necessary for mouse preimplantation embryo development and cell-fate specification. Open Biol 11: 210092.

Bora P, Gahurova L, Masek T, Hauserova A, Potesil D, Jansova D, Susor A, Zdrahal Z, Ajduk A, Pospisek M et al. 2021b. p38-MAPK-mediated translation regulation during early blastocyst development is required for primitive endoderm differentiation in mice. Commun Biol 4: 788.

Bora P, Thamodaran V, Susor A, Bruce AW. 2019. p38-Mitogen Activated Kinases Mediate a Developmental Regulatory Response to Amino Acid Depletion and Associated Oxidative Stress in Mouse Blastocyst Embryos. Front Cell Dev Biol 7: 276.

Chazaud C, Yamanaka Y. 2016. Lineage specification in the mouse preimplantation embryo. Development 143: 1063–1074.

Chazaud C, Yamanaka Y, Pawson T, Rossant J. 2006. Early lineage segregation between epiblast and primitive endoderm in mouse blastocysts through the Grb2-MAPK pathway. Dev Cell 10: 615–624.

Del Llano E, Masek T, Gahurova L, Pospisek M, Koncicka M, Jindrova A, Jansova D, Iyyappan R, Roucova K, Bruce AW et al. 2020. Age-related differences in the translational landscape of mammalian oocytes. Aging Cell 19: e13231.

Fischer SC, Corujo-Simon E, Lilao-Garzon J, Stelzer EHK, Munoz-Descalzo S. 2020. The transition from local to global patterns governs the differentiation of mouse blastocysts. PLoS One 15: e0233030.

Fonseca BD, Smith EM, Yelle N, Alain T, Bushell M, Pause A. 2014. The ever-evolving role of mTOR in translation. Semin Cell Dev Biol 36: 102–112.

Frankenberg S, Gerbe F, Bessonnard S, Belville C, Pouchin P, Bardot O, Chazaud C. 2011. Primitive endoderm differentiates via a three-step mechanism involving Nanog and RTK signaling. Dev Cell 21: 1005–1013.

Gahurova L, Tomankova J, Cerna P, Bora P, Kubickova M, Virnicchi G, Kovacovicova K, Potesil D, Hruska P, Zdrahal Z et al. 2023. Spatial positioning of preimplantation mouse embryo cells is regulated by mTORC1 and m(7)G-cap-dependent translation at the 8- to 16-cell transition. Open Biol 13: 230081.

Iyyappan R, Aleshkina D, Ming H, Dvoran M, Kakavand K, Jansova D, Del Llano E, Gahurova L, Bruce AW, Masek T et al. 2023. The translational oscillation in oocyte and early embryo development. Nucleic Acids Res 51: 12076–12091.

Jackson JR, Bolognese B, Hillegass L, Kassis S, Adams J, Griswold DE, Winkler JD. 1998. Pharmacological effects of SB 220025, a selective inhibitor of P38 mitogen-activated protein kinase, in angiogenesis and chronic inflammatory disease models. J Pharmacol Exp Ther 284: 687–692.

Kalmar T, Lim C, Hayward P, Munoz-Descalzo S, Nichols J, Garcia-Ojalvo J, Martinez Arias A. 2009. Regulated fluctuations in nanog expression mediate cell fate decisions in embryonic stem cells. PLoS Biol 7: e1000149.

Kang M, Piliszek A, Artus J, Hadjantonakis AK. 2013. FGF4 is required for lineage restriction and salt-and-pepper distribution of primitive endoderm factors but not their initial expression in the mouse. Development 140: 267–279.

Lou X, Kang M, Xenopoulos P, Munoz-Descalzo S, Hadjantonakis AK. 2014. A rapid and efficient 2D/3D nuclear segmentation method for analysis of early mouse embryo and stem cell image data. Stem Cell Reports 2: 382–397.

Manejwala FM, Cragoe EJ, Jr., Schultz RM. 1989. Blastocoel expansion in the preimplantation mouse embryo: role of extracellular sodium and chloride and possible apical routes of their entry. Dev Biol 133: 210–220.

Masek T, Del Llano E, Gahurova L, Kubelka M, Susor A, Roucova K, Lin CJ, Bruce AW, Pospisek M. 2020. Identifying the Translatome of Mouse NEBD-Stage Oocytes via SSP-Profiling; A Novel Polysome Fractionation Method. Int J Mol Sci 21.

Mathew B, Munoz-Descalzo S, Corujo-Simon E, Schroter C, Stelzer EHK, Fischer SC. 2019. Mouse ICM Organoids Reveal Three-Dimensional Cell Fate Clustering. Biophys J 116: 127–141.

McCloy RA, Rogers S, Caldon CE, Lorca T, Castro A, Burgess A. 2014. Partial inhibition of Cdk1 in G 2 phase overrides the SAC and decouples mitotic events. Cell Cycle 13: 1400–1412.

Mihajlovic AI, Bruce AW. 2016. Rho-associated protein kinase regulates subcellular localisation of Angiomotin and Hippo-signalling during preimplantation mouse embryo development. Reprod Biomed Online 33: 381–390.

Mihajlovic AI, Thamodaran V, Bruce AW. 2015. The first two cell-fate decisions of preimplantation mouse embryo development are not functionally independent. Sci Rep 5: 15034.

Motosugi N, Bauer T, Polanski Z, Solter D, Hiiragi T. 2005. Polarity of the mouse embryo is established at blastocyst and is not prepatterned. Genes Dev 19: 1081–1092.

Niakan KK, Ji H, Maehr R, Vokes SA, Rodolfa KT, Sherwood RI, Yamaki M, Dimos JT, Chen AE, Melton DA et al. 2010. Sox17 promotes differentiation in mouse embryonic stem cells by directly regulating extraembryonic gene expression and indirectly antagonizing self-renewal. Genes Dev 24: 312–326.

Nichols J, Silva J, Roode M, Smith A. 2009. Suppression of Erk signalling promotes ground state pluripotency in the mouse embryo. Development 136: 3215–3222.

Ozadam H, Tonn T, Han CM, Segura A, Hoskins I, Rao S, Ghatpande V, Tran D, Catoe D, Salit M et al. 2023. Single-cell quantification of ribosome occupancy in early mouse development. Nature 618: 1057–1064.

Plusa B, Piliszek A. 2020. Common principles of early mammalian embryo self-organisation. Development 147.

Posfai E, Petropoulos S, de Barros FRO, Schell JP, Jurisica I, Sandberg R, Lanner F, Rossant J. 2017. Position- and Hippo signaling-dependent plasticity during lineage segregation in the early mouse embryo. Elife 6.

Potapova TA, Sivakumar S, Flynn JN, Li R, Gorbsky GJ. 2011. Mitotic progression becomes irreversible in prometaphase and collapses when Wee1 and Cdc25 are inhibited. Mol Biol Cell 22: 1191–1206.

Ryan AQ, Chan CJ, Graner F, Hiiragi T. 2019. Lumen Expansion Facilitates Epiblast-Primitive Endoderm Fate Specification during Mouse Blastocyst Formation. Dev Cell 51: 684–697 e684.

Schindelin J, Arganda-Carreras I, Frise E, Kaynig V, Longair M, Pietzsch T, Preibisch S, Rueden C, Saalfeld S, Schmid B et al. 2012. Fiji: an open-source platform for biological-image analysis. Nat Methods 9: 676-682.

Schliffka MF, Dumortier JG, Pelzer D, Mukherjee A, Maître J-L. 2023. Inverse blebs operate as hydraulic pumps during mouse blastocyst formation. bioRxiv doi:10.1101/2023.05.03.539105:2023.2005.2003.539105.

Schroter C, Rue P, Mackenzie JP, Martinez Arias A. 2015. FGF/MAPK signaling sets the switching threshold of a bistable circuit controlling cell fate decisions in embryonic stem cells. Development 142: 4205–4216.

Solter D, Knowles BB. 1975. Immunosurgery of mouse blastocyst. Proc Natl Acad Sci U S A 72: 5099–5102.

Thamodaran V, Bruce AW. 2016. p38 (Mapk14/11) occupies a regulatory node governing entry into primitive endoderm differentiation during preimplantation mouse embryo development. Open Biol 6.

Watson AJ. 1992. The cell biology of blastocyst development. Mol Reprod Dev 33: 492–504.

Watson AJ, Barcroft LC. 2001. Regulation of blastocyst formation. Front Biosci 6: D708–730.

Wicklow E, Blij S, Frum T, Hirate Y, Lang RA, Sasaki H, Ralston A. 2014. HIPPO Pathway Members Restrict SOX2 to the Inner Cell Mass Where It Promotes ICM Fates in the Mouse Blastocyst. PLoS Genet 10: e1004618.

Wigger M, Kisielewska K, Filimonow K, Plusa B, Maleszewski M, Suwinska A. 2017. Plasticity of the inner cell mass in mouse blastocyst is restricted by the activity of FGF/MAPK pathway. Sci Rep 7: 15136.

Wiley LM. 1984. Cavitation in the mouse preimplantation embryo: Na/K-ATPase and the origin of nascent blastocoele fluid. Dev Biol 105: 330–342.

Yamanaka Y, Lanner F, Rossant J. 2010. FGF signal-dependent segregation of primitive endoderm and epiblast in the mouse blastocyst. Development 137: 715–724.

